# Accurate Simulation of Coupling between Protein Secondary Structure and Liquid-Liquid Phase Separation

**DOI:** 10.1101/2023.08.22.554378

**Authors:** Yumeng Zhang, Shanlong Li, Xiping Gong, Jianhan Chen

## Abstract

Intrinsically disordered proteins (IDPs) frequently mediate liquid-liquid phase separation (LLPS) that underlies the formation of membraneless organelles. Together with theory and experiment, efficient coarse-grained (CG) simulations have been instrumental in understanding sequence-specific phase separation of IDPs. However, the widely-used Cα-only models are severely limited in capturing the peptide nature of IDPs, including backbone-mediated interactions and effects of secondary structures, in LLPS. Here, we describe a hybrid resolution (HyRes) protein model for accurate description of the backbone and transient secondary structures in LLPS. With an atomistic backbone and coarse-grained side chains, HyRes accurately predicts the residue helical propensity and chain dimension of monomeric IDPs. Using GY-23 as a model system, we show that HyRes is efficient enough for direct simulation of spontaneous phase separation, and at the same time accurate enough to resolve the effects of single mutations. HyRes simulations also successfully predict increased beta-sheet formation in the condensate, consistent with available experimental data. We further utilize HyRes to study the phase separation of TPD-43, where several disease-related mutants in the conserved region (CR) have been shown to affect residual helicities and modulate LLPS propensity. The simulations successfully recapitulate the effect of these mutants on the helicity and LLPS propensity of TDP-43 CR. Analyses reveal that the balance between backbone and sidechain-mediated interactions, but not helicity itself, actually determines LLPS propensity. We believe that the HyRes model represents an important advance in the molecular simulation of LLPS and will help elucidate the coupling between IDP transient secondary structures and phase separation.

## Introduction

Membraneless organelles or biological condensates are now recognized to be an important mechanism of subcellular compartmentalization and contribute to myriad biological functions such as RNA storage and processing, stress response, cellular trafficking, metabolism, and cellular signaling.^1–4^ They have also been linked to many diseases including cancer, neurodegeneration, and infectious diseases.^5–10^ Extensive studies in recent years have established that biological condensates are viscous liquid-like structures, which form and dissolve through spontaneous phase separations driven by dynamic and multivalent biomacromolecules such as RNAs and proteins.^11–13^ One of the major drivers of the initial liquid-liquid phase separation (LLPS) is intrinsically disordered proteins or regions (IDPs/IDRs).^14–17^ IDPs are enriched with charged and polar residues, have lower sequence complexity, lack stable tertiary structures, and are key components of cellular signaling and regulatory networks.^18–23^ The dynamic and multivalent nature allows IDPs to frequently serve as scaffolds of biological condensates.^3, 24^ Maturation frequently follows LLPS of IDPs, where the droplets become increasingly viscoelastic and turn into fibrils, gels, glasses, or other solid-like materials.^2^ Intensive efforts have been dedicated to understanding how the balance of various solvent and co-solvent mediated backbone and side chain interactions together give rise to a rich spectrum of sequence-specific phase separation properties of IDPs, by themselves or together with RNAs and other molecules.^25–27^

Classical concepts from polymer physics have been central to understanding the general physical principles governing the biological condensates of IDPs.^1, 28–31^ For example, the Flory-Huggins mean-field formalism for homopolymers captures the balance of polymer-solvent and polymer-polymer interactions with a single Flory-Huggins *χ* parameter,^32, 33^ and can be effectively used to fit and understand the phase diagram of IDPs.^34–36^ Random phase approximations (RPAs) can account for electrostatic effects on the phase transitions of polyampholytes with any charge pattern, allowing the prediction of sequence and pH/salt dependence of IDP phase separation.^37,38^ Field-theoretic simulations (FTS) have been further developed to numerically sample the fully fluctuating field-theoretic Hamiltonian and alleviate the restriction of analytical approximations in RPAs.^39–41^ FTS approaches enable more realistic descriptions of the roles of charge-charge interactions as well as sequence-dependent short- and long-range interactions in driving phase separations of IDPs. The stickers-and-spacers models for associative polymers provide a particularly effective framework for coarse-grained (CG) lattice simulations of the phase transitions of multivalent proteins to decipher the impacts of various sequence features.^26, 42, 43^ These models are applicable to multi-component systems and can describe additional structural features such as percolation and gelation in dilute and condensed phases.

Compared to theoretical approaches, molecular dynamics (MD) simulations using transferable energy functions can be particularly attractive for deriving system-specific molecular details of phase separation, because they are free of simplifying assumptions required for making the theories tractable.^44, 45^ In particular, CG protein models are often necessary to access the large length (> ∼100 nm) and time scales (> μs) required for direct simulation of IDP phase separation. Among them, Cα-only models have been the most successful in simulating LLPS of IDPs. Here, each residue is represented by a single “Cα” CG bead, interacting through effective potentials derived from structural statistics, atomistic simulations, and/or experimental observables. Similar Cα-only models, such as Gō-like models, have been quite successful in studies of protein folding,^46–49^ transient protein-protein interactions,^50, 51^ and coupled binding and folding of IDPs.^52–55^ The original Kim-Hummer (KH) model was first adopted by Mittal, Best, and coworkers for simulating IDP phase separations,^56^ where an alternative version with the original Miyazawa-Jerningan (MJ) statistical contact potential replaced by a hydrophobicity scale (HPS)-based one was also explored. Both KH and HPS models were shown to be capable of providing qualitative prediction of changes in the phase diagram of low complexity domain (LCD) of protein FUS due to mutations or the presence of folded protein LAF-1. Many variants of Cα-only models have since been developed with contact pair potentials assigned and optimized using different strategies including machine learning and sometimes with attention to specific interaction types such as π-π contacts^57–66^. Together, these models have demonstrated increasingly robust abilities to recapitulate experimental observables such as monomer radius of gyration (*R*_g_) and phase transition boundaries. Multi-bead CG protein models have also been explored for simulation of IDP phase separation.^67–69^ For example, the popular MARTINI force field can be rebalanced to simulate the droplet formation of FUS LCD.^69^ A range of liquid properties including densities, surface tensions, and sheer viscosities could be reproduced with moderate downscaling (∼0.65) of the original MARTINI protein-protein interactions.

A critical limitation of Cα-only or MARTINI models, however, is their inability to accurately describe the interactions and secondary structures of the peptide backbone, due to the single bead representation of the entire residue (Cα-only models) or backbone (MARTINI). Yet, it is well recognized that anisotropic backbone interactions are important in driving IDP phase separations.^1, 44, 70–72^ Critically, IDPs are not simple polymers, and transient secondary and sometimes long-range structures are prevalent.^73, 74^ Recent studies have highlighted the strong interplay between transient secondary structures and phase separation behaviors.^75–78^ For instance, IDPs and IDRs often adopt more extended conformations within condensates and often with an increased propensity for *β*-structures.^79–81^ It has also suggested LLPS can induce the folding of poly-(PR) peptides into helical conformations,^77, 82^ which is a characteristic feature of aging-related phase transition pathologies. Furthermore, recent studies have revealed that the C-terminal domain of TAR DNA-binding protein 43 (TDP-43) undergoes transient conformational changes during phase transitions, forming helices during LLPS and subsequently aging into amyloid structures.^83^ Intriguingly, TDP-43 mutants with enhanced helicity exhibit a higher LLPS propensity, suggesting a driving role of residual helices in TDP-43 phase separation.^84, 85^

Accurate description of the peptide backbone and transient secondary structures arguably requires near atomistic representation beyond the single bead Cα-only resolution.^86^ This is because backbone interactions are highly anisotropic in nature, and the anisotropic nature is responsible for the conformational dependence of these interactions. For example, the backbone within a helix segment is no longer available to form hydrogen bonding with water molecules or the rest of the protein. This conformational dependence of backbone interactions cannot be captured using Cα-only models, even if empirical potentials are introduced to mimic sequence-dependent helical structures.^87^ At present, atomistic simulations largely remain the only option for examining details of peptide backbone interactions and potentially the effects of transient structures in LLPS of IDPs.^72, 88–91^ While these courageous attempts have provided useful insights, atomistic simulations of LLPS are severely limited by the achievable simulation timescales, and deriving statistically meaningful predictions of biological condensates is largely out of reach. There is a critical need for simplified protein models that accurately describe the peptide backbone and at the same time are computationally trackable for simulation of IDP phase separation.

We recently developed a hybrid resolution (HyRes) protein model for accurate description of local and long-range structures of IDPs.^92, 93^ As illustrated in Figure 1A, the HyRes model contains fully atomistic backbones, and the side chains are represented by up to five CG beads (for Trp). The energy function of HyRes contains bond, angle, dihedral/improper, backbone CMAP dihedral cross-term, Lennard-Jones (LJ) van der Waals (vdW), Debye-Hückel electrostatics, and backbone hydrogen bonding and solvent accessible surface area (SASA)-based implicit solvent. The bonded terms as well as the LJ radii were parameterized to reproduce the side chain geometry, volume, and flexibility from atomistic simulations. The relative strengths of LJ potentials (ε_i_) for all side chains CG beads were first assigned based on the MJ statistical contact potentials,^94^ and then uniformly rescaled to balance electrostatics and hydrogen-bonding terms based on a set of small model peptides. The latest version, HyRes II, has been shown to provide nearly quantitative description of a range of nontrivial local and long-range structure features of IDPs, including residual helicity and transient long-range contacts, at a level comparable to atomistic simulations.^93^ It was also able to describe the dynamic interaction between the N-terminal transactivation domain (TAD) of tumor suppressor p53 and protein cyclophilin D, predicting structure features that are highly consistent with NMR titration and binding experiments.^95^ Therefore, HyRes may provide an appropriate CG protein model for direct simulation of peptide backbone and secondary structures in LLPS of IDPs.

**Figure 1.**
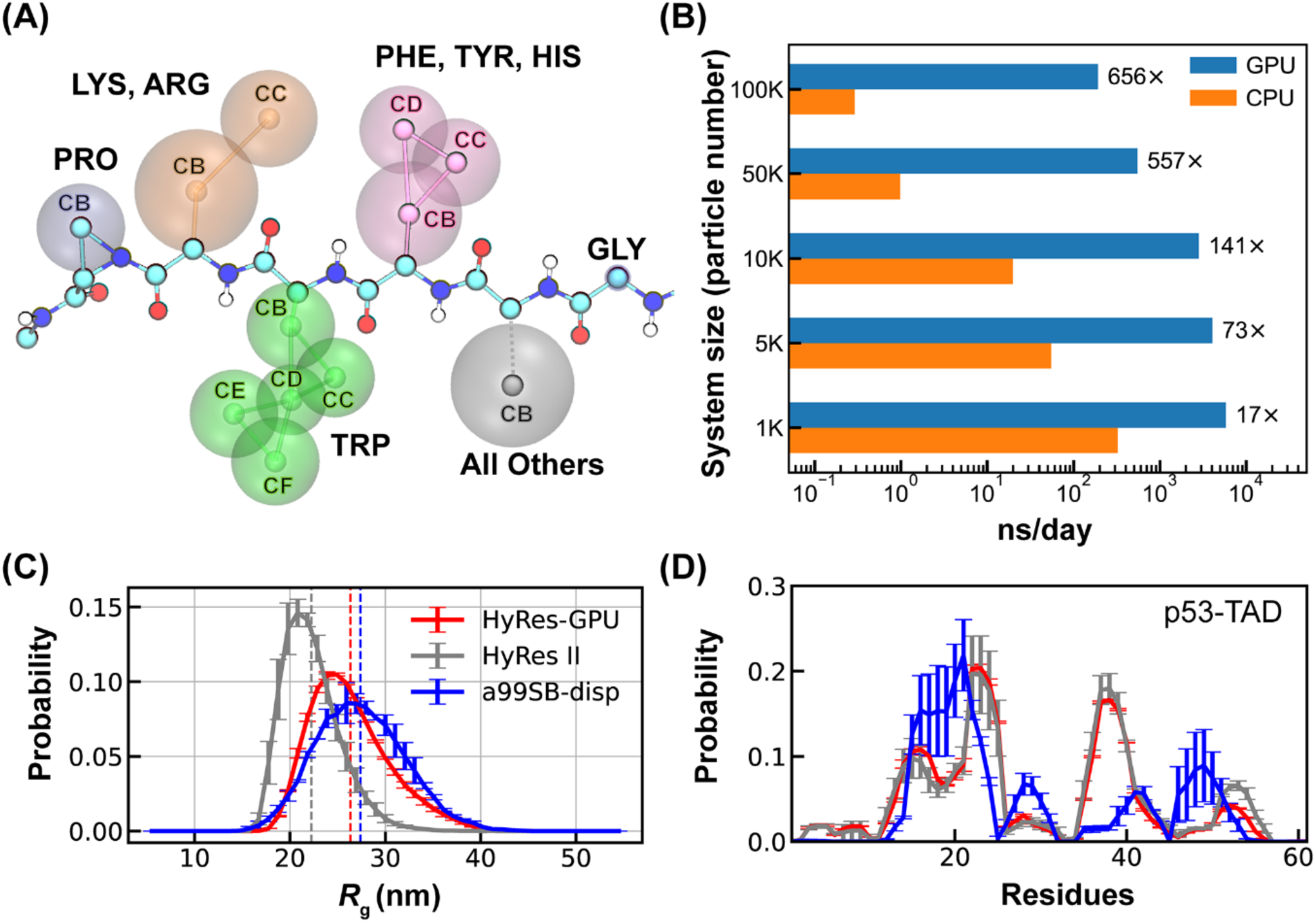
Representation, efficiency, and accuracy of HyRes-GPU. (A) Physical representation of the HyRes model. The backbone atoms are drawn in CPK style with carbon in cyan, nitrogen in blue, oxygen in red, and hydrogen in white. The side-chain beads are illustrated by vdW spheres. (B) Computational efficiencies of HyRes-GPU in comparison to HyRes II for systems of different sizes. The timing simulations were performed using either a single Xeon E5-2620 CPU core or a single Nvidia RTX 4070Ti GPU card. (C-D) Probability distributions of *R*_g_ and residue helicity profiles of p53-TAD (1-61) simulated using HyRes-GPU, in comparison to results from HyRes II (grey traces) and atomistic force field a99SB-disp (blue traces). The vertical dashed lines mark average *R*_g_ values of the simulated ensembles. The atomistic results are taken from a previous work.^101^

Here, we develop a GPU-accelerated HyRes model, and critically assess its suitability for simulating spontaneous phase separation of IDPs and studying the roles of backbone-mediated interactions and residual structures in LLPS. We show that the HyRes-GPU model is able to directly simulation LLPS of GY-23 peptides derived from histidine-rich beak proteins (HBPs) and recapitulate the effects of single mutations on the phase separation propensity. The simulations also correctly predict increased *β*-structures in the condensed phase, consistent with previous CD measurements. We further examine the molecular basis of how residual helicity of the conserved region (CR) of TDP-43 modulates its phase separation propensity. HyRes simulations correctly predict a positive correlation between monomeric peptide helicity and condensate stability as measured by the saturation concentration (*C*_sat_). However, analyses based on HyRes simulations reveal that residual helices do not play a direct role by increasing helix-helix interactions in the condensed phase as previously proposed. Instead, it is the balance among solvation, side chain hydrophobicity, and availability of backbone that actually governs the condensate stability of TDP- 43 CR. These results suggest that HyRes provides an efficient and viable model for simulating IDP phase separation and interrogating the role of peptide backbone and residual structures.

## Results and Discussion

### GPU acceleration of the HyRes protein model

We first implemented the HyRes model on GPU within the OpenMM package^96–98^ to dramatically expand the accessible length- and time-scales required for studying IDP phase separation. The final HyRes-GPU implementation can provide several hundred-fold speed-ups compared to the CPU version for large systems (Figure 1B). The production rate is near 200 ns/day for a 100 K atoms system on a single Nvidia GTX 4070Ti card. We note that the SASA solvation energy term, applied only to the backbone, is removed in HyRes-GPU and compensated by recalibrating the backbone LJ parameters for computational efficiency (Table S1). The recalibration was guided by examining conformational properties of two model IDPs, (AAQAA)_3_ and KID (see Table S2 for sequences). The new parameters for backbone atoms, summarized in Table S1, allow HyRes- GPU to essentially reproduce both residual helical structures and overall compaction of both peptides (Figure S1). We further validated the balance of HyRes-GPU using p53-TAD (residue 1- 61), a longer IDP with nontrivial structural properties (Figure 1C-D). The result shows that HyRes- GPU fully retains the ability of HyRes II in predicting residual helical propensities. In particular, HyRes-GPU provides further improvement in capturing the level of overall compaction as reflected in the distribution of radius of gyration (*R*_g_), compared well to the result from a99SB-disp, one of the best explicit solvent atomistic protein force fields.^99, 100^

### Ability of HyRes-GPU to describe properties of monomeric IDPs

It has been shown that single-molecule properties of IDPs are intimately connected to their phase behavior.^102, 103^ For example, the coil-to-globule transition temperature can be used as a proxy for predicting the upper critical solution temperature.^102^ In fact, single-chain properties, particularly *R*_g_, have been frequently used in the optimization of CG protein models for the simulation of LLPS.^64–66^ Models better reproducing experimental single-chain *R*_g_ have been shown to more accurately predict homotypic LLPS propensities. Therefore, we first examine the ability of HyRes- GPU to reproduce the experimental *R*_g_ of a set of IDPs ranging from 24 to 236 residues in length (Table S2). Some of these IDPs have been shown to drive LLPS, while others have not. The test set also exhibits significant variation in charge distribution (from -44 to +16), which can help to evaluate balance among local contacts, long-range nonpolar and electrostatic interactions in the model. As summarized in Figure 2A, *R*_g_ values computed from HyRes-GPU simulations agree well with the experimental results, with a Pearson’s correlation score of *r* = 0.66. This compares well to early Cα-only models such as KH (*r* ∼0.54)^64^ and HPS or HPS-Urry (*r* ∼0.3 – 0.6).^65^ This is notable because HyRes-GPU has not been directly optimized to reproduce experimental *R*_g_ of a large number of IDPs. Instead, HyRes has only been parameterized based on small model peptides including poly-Gly, (AAQAA)_3_, KID and the (EK)_25_ series, to achieve quantitative long-range non-specific interactions and qualitative residual structure descriptions.^92, 93^ Along this line, it is unsurprising that HyRes-GPU yields lower Pearson’s correlation score compared to *R*_g_- optimized models, such as Mpipi,^64^ HPS-cation-π^61^ and others.^65, 66^ The implication is also that HyRes-GPU could be further improved following similar optimization strategies.

**Figure 2.**
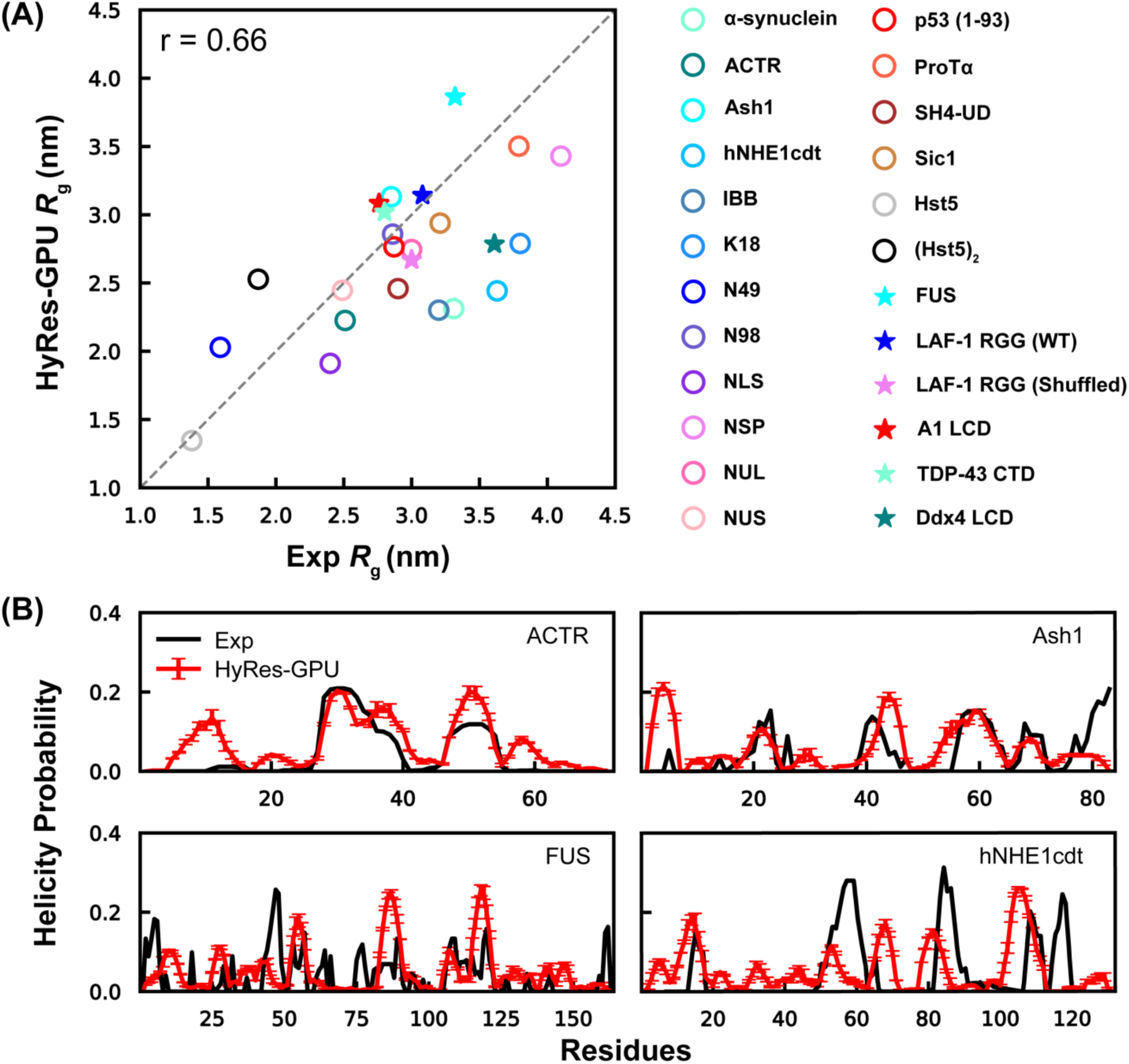
Monomeric conformational properties of IDPs in HyRes-GPU. (A) Correlation between *R*_g_ from simulation and experiment for a set of IDPs. The sequences are provided in Table S2. (B) Residue helicity profiles for four representative IDPs from HyRes-GPU, in comparison to experimental results for ACTR,^104^ Ash1,^105^ FUS,^106^ and hNHE1cdt.^107^

We further analyzed the residual structures of several IDPs known to exhibit dynamic local conformations with available experimental data (Figure 2B). The results support that HyRes-GPU can at least semi-quantitatively predict sequence-specific helical propensities. Although there are some mismatched local structures that are over- or under-estimated, residual helices within the structural ensembles generated by HyRes-GPU are generally consistent with experiments. For example, the model accurately predicts ∼20% helicity between 25-40 residues of ACTR, ∼15 % helicity between 55-65 residues of Ash1, and the helicity between residues 105-118 of FUS. Particularly, hNHE1cdt is not overly helical even though its chain dimension is slightly underestimated by HyRes-GPU (as indicated by its *R*_g_ values in Figure 2A).

### HyRes-GPU simulation of spontaneous phase separation of GY-23

To examine the ability of HyRes-GPU for simulating LLPS, we first focus on a short 23-residue low complexity peptide GY-23 (see Table S2 for sequence), the central segment of squid-beak derived protein HBP-1 that regulates its pH-dependent LLPS propensity.^72^ GY-23 alone can undergo homotypic LLPS at room temperature and neutral pH of 7.0. Importantly, its phase behavior is highly sensitive to even single-point mutations.^108^ Therefore, GY-23 provides a good model system for evaluating the suitability of HyRes-GPU for direct simulation of LLPS and further examining if the model is accurate enough to recapitulate the effects of single-point mutations. For this, we performed multiple simulations of 200 copies of wild-type (WT) GY-23 in a 45-nm cubic box for a total effective concentration of ∼3.6 mM, starting from either preformed high-density compact or fully dispersed initial states at 300 K (see Methods and SI Figure S2 for the generation of initial configurations). As illustrated in Figure 3, the system reached similar phase separated states within ∼1 *µs* regardless of the initial configurations. Representative movies of the simulation trajectories are provided in the Supplementary Materials (Movies S1 and S2). A single dominant droplet was observed in the second half of all six independent simulations. The droplets are of comparable sizes (Figure 3C) and have largely identical density profiles (Figures S3 and 3D). The droplet formation from the dispersed initial configuration apparently follows the classical homogeneous nucleation mechanism, where small clusters would form and dissolve during the initial stages of phase separation (Figure 3C). Once exceeding the critical nucleus size, the droplet would grow rapidly until reaching the final equilibrium. We further examine the finite size effects on the droplet by simulating the phase equilibrium with 100, 200, and 300 copies in three different box sizes at 300 K. The results, summarized in Figure S4, show that the droplets have similar density profiles and reach the same high concentration in the droplet interior in all three cases. As such, all additional simulations of WT GY-23 and mutants at various temperatures were performed using 200 copies in the 45-nm cubic box.

**Figure 3.**
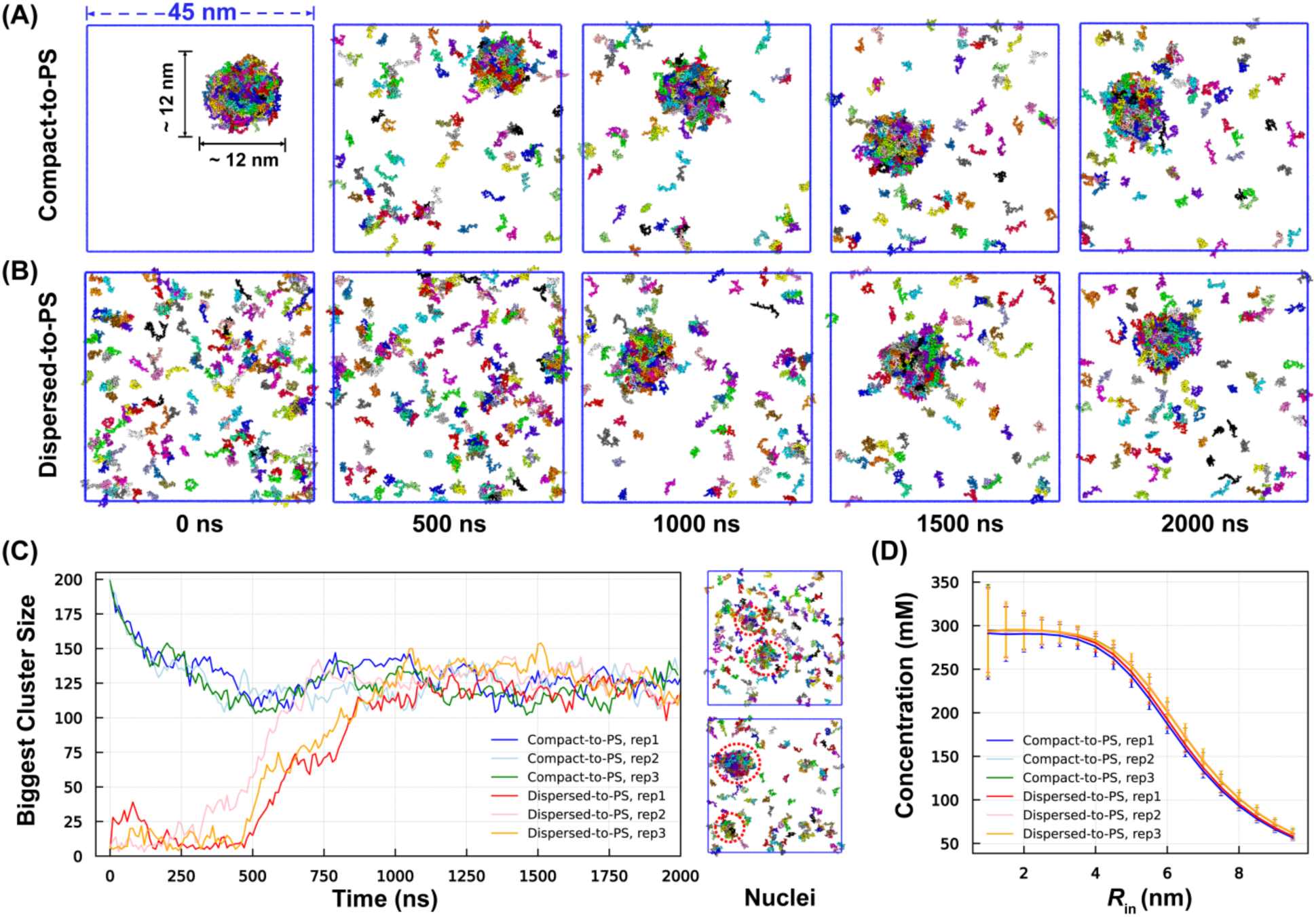
Spontaneous phase separation of GY-23 in HyRes-GPU. (A, B) Snapshots from representative simulations initiated from either compact or dispersed initial states, showing the system reaching a similar phase separated (PS) final state during the course of 2 *µs* simulation timescale. (C) The size of the largest cluster during three replicas of simulations initiated from either the compact or dispersed initial states. The panels illustrate that multiple small nuclei (red circles) often form and dissolve during the initial stage of simulations from the dispersed state. (D) The concentration profiles of the final condensed phase were derived from all 6 simulations. The error bars were estimated from standard deviation of five block averages within the last 500 ns trajectories.

We note that the peptides undergo rapid exchange between the condensed and dilute phases throughout the HyRes-GPU simulations. Figure S5A plots the numbers of GY-23 peptides that never leaves the droplet as a function of time from representative trajectories of simulations initiated from the compact configuration at different temperatures. It shows that the number of peptides that had yet exchanged with the dilute phase decreased rapidly to zero, within ∼250 ns at 300 K, even though the equilibrium droplet size is ∼125 (Figure 3C). The rapid dynamics of peptide exchange between two phases in HyRes-GPU is a highly desirable property; it allows efficient simulation of the dynamic phase equilibrium to derive reliable thermodynamic properties and potentially resolve the effects of mutations and other perturbations. We further characterize the peptide diffusion constants of dilute and condensed phases by analyzing the mean-squared displacement (MSD) functions (Figure S5B). The results show that the diffusion constant of GY- 23 is about 10-fold slower in the droplet compared to the dilute phase. This is smaller than the ∼100-fold experimental estimate.^109^ Multiple factors may contribute to the underestimation of viscosity increase in the condensate, including smoother effective potentials used in CG models such as HyRes-GPU. In addition, the size of the droplet simulated here (∼10 nm in diameter) is too small to capture longer range ordering that gives rise to slower diffusion kinetics and viscoelastic material properties.^28, 110^

### HyRes-GPU recapitulates mutational effects on GY-23 phase equilibrium

It has been shown experimentally that the ability of GY-23 to phase separate is sensitive to Histidine-to-Lysine (H-to-K) mutations.^108^ Either single H-to-K mutations, H2K, H12K, H15K, and H20K, or all H-to-K mutations, H/K, can significantly shift the phase co-existence, such that phase separation is completely abolished for H2K, H20K, and H/K mutants at all tested conditions and can only be achieved at higher pH (> 8) and/or salt concentrations (0.5 or 1 M NaCl) for H12K and H15K mutants. A set of simulations at different temperatures spanning 260 to up to 310 K were performed for each GY-23 construct (including WT) to probe the impact of mutations on phase equilibrium. All simulations were initiated from a preformed high-density compact configuration (see Method and Figure S4). The results show that, with a total concentration of ∼3.6 mM, the preformed droplet was quickly dissolved for all H-to-K mutants at 300 K (Figure S6) and there is no phase separation at the end of the 2-*µs* simulations unlike the case of WT GY-23 (Figure 4A). This is apparently consistent with the experimental observations.^108^ Interestingly, whereas the H/K mutant shows no phase separation ability even at 260 K, the single H-to-K mutants can undergo phase separation at lower temperatures. To further quantify the impacts of H-to-K mutations on phase separation of GY-23, we constructed the phase co-existence curves and derived the critical phase separation temperature (*T*_c_) and concentration (*ρ*_c_) (Figure 4B). The H/K mutant was not included in the diagram due to the absence of phase separation at all temperatures simulated. The critical concentration for phase separation of the WT peptide is ∼1.5 mM, consistent with the experimental result showing that phase separation occurs at 0.5 mM or higher.^108^ For the single H-to-K mutants, the critical concentration of phase separation is >6 mM at 300 K, which is consistent with a lack of phase separation experimentally examined between 0.1 to 1.0 mM. Furthermore, fitting the phase coexistence curves to Equations 1 and 2 shows a significant decrease in *T*_c_ for all single H-to-K mutants. The ability of HyRes-GPU to recapitulate the impact of single H-to-K mutations on the phase behavior of GY-23 is encouraging. It suggests that the balance between various interactions, particularly nonpolar vs. electrostatic, is well captured in HyRes-GPU, which is crucial for accurate simulation of how sequence and other structural properties of IDPs may modulate phase separation.

**Figure 4.**
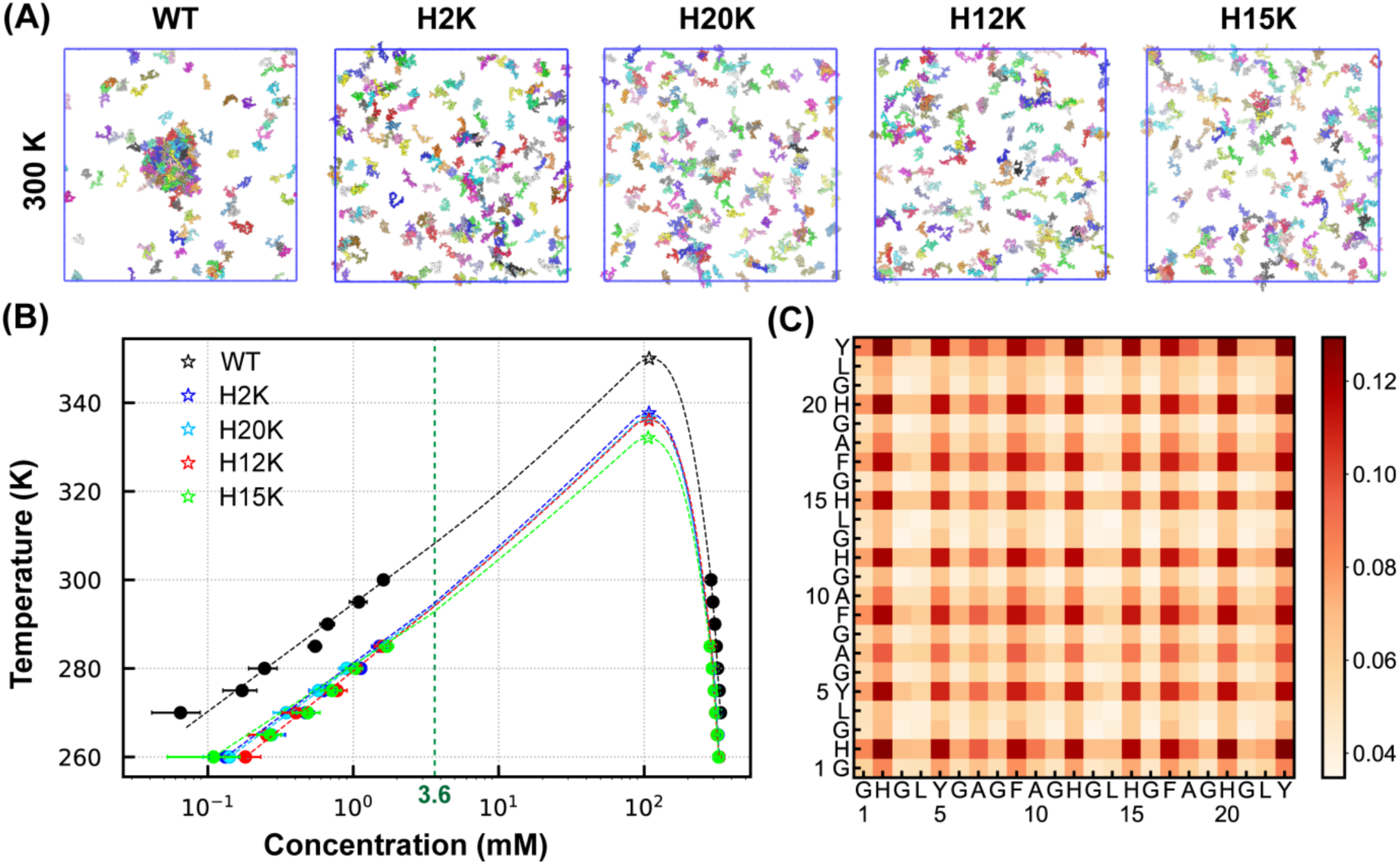
Phase separation of GY-23 peptide variants. (A) Representative final snapshots of the 2-*µs* simulations of WT GY-23 and four single H-to-K mutants in HyRes-GPU at 300 K. (B) Phase diagram of GY-23 WT and single H-to-K mutants. The error bars shown were estimated from standard deviation of three block averages within the last 500 ns trajectories. *T*_c_ and *ρ*_c_ (marked as “☆”) were estimated by fitting to Equations 1 and 2 (see Methods). Dashed smooth curves were drawn to show the trend of phase boundaries. The total peptide concentration is ∼3.6 mM in all simulations, marked by the green dashed vertical line. (C) Probabilities of intermolecular residue-residue contacts in the condensed phase of WT GY-23 at 300 K.

### Molecular basis of GY-23 phase separation: importance of aromatic residues and coupling with *β*-structure formation

To further elucidate the molecular basis of GY-23 phase separation, we first analyze the probabilities of intermolecular contacts in the condensate and compare the peptide conformational properties in the dilute and condensed phases. The results reveal that there is only a slight increase in the peptide chain expansion upon forming the condensate, as reflected in the distributions of *R*_g_ and end-to-end distance (*R*_ee_) (Figure 5A and B). This is not surprising because GY-23 is highly disordered in both phases. Monomeric GY-23 peptides display minimal secondary structures at 300 K (∼1%, Figure S7), which is consistent with previous CD measurements.^109^ The propensity for greater extension in the condensed phase has been observed for IDPs in numerous other cases.^111–114^

**Figure 5.**
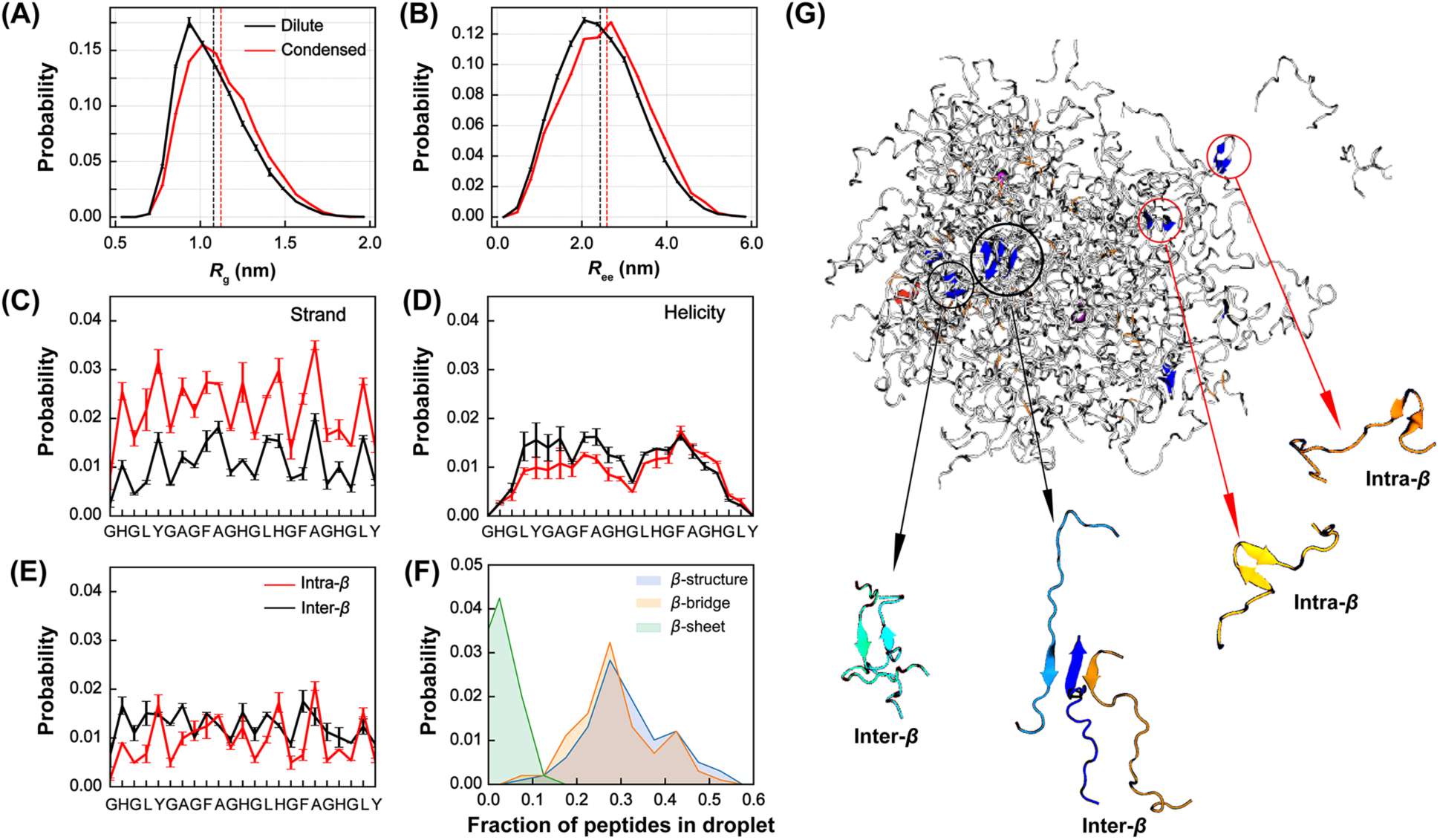
Conformational properties of WT GY-23 at phase coexistence at 300 K. (A-D) Distributions of *R*_g_, *R*_ee_, residue *β*-structure propensity and helicity in dilute (black) and condensed (red) phases. (D) Probability of each residue forming intra- vs inter-peptide *β*-sheet structures in the condensed phase. (F) Probability distributions of the fraction of peptides that contain any *β*- structures (blue), participate in *β*-bridges (green), or *β*-sheets (cyan) in the condensed phase. (G) A representative snapshot of the droplet with *β*-sheet and helical structures colored in blue and red, respectively. Note that ∼1/3 of peptides are involved in *β*-bridges and they are not highlighted.

A notable difference in the GY-23 conformations in dilute and condensed phases is a significant increase in the propensity for *β*-structures within the condensed phase (Figure 5C). In contrast, the peptide contains similarly low helical content in both phases (Figure 5D). Although the overall probability of forming *β*-strands and sheets remains low (∼3% per residue), the presence of *β*- structures (including *β*-bridges) within the condensate likely plays a significant role in driving phase separation. In particular, within the droplet at 300 K, even though only ∼5% of the peptides participate in intermolecular parallel or anti-parallel *β*-sheets, almost 35% of the peptides are involved in *β*-bridges and over half of these are intermolecular ones (Figure 5E and F). A representative snapshot of the equilibrium droplet at 300 K is shown in Figure 5G, where the presence of several intra- and intermolecular *β*-sheets are highlighted. We note that the level of intermolecular *β*-structures increases dramatically and faster than intramolecular ones at lower temperatures (Figure S8). At 260 K, over 50% of peptides are involved in *β*-structures, of which over 70% are intermolecular ones, apparently contributing to more stable phase separation. The importance of *β*-structures, and particularly an apparent coupling between *β*-structures and condensate formation, has also been suggested in a previous experimental study of a longer 161- residue HBP-1 construct.^109^ CD spectra reveal a negative peak at 215 - 220 nm upon condensation formation at high HBP-1 concentration, consistent with a shift towards higher *β*- structure. These results strongly support a coupling between protein secondary structures and phase separation and highlight the importance of accurate description of protein backbone and transient secondary structures.

We further examine the molecular basis of how H-to-K mutations affect the ability of GY-23 to phase separate. Analysis of the monomeric conformational properties, summarized in Figure S7, shows that single H-to-K mutations have minimal impacts on compaction or secondary structural propensities. This is not surprising given the highly unstructured nature of GY-23. Only replacing all four histidines with lysines in the H/K mutant can lead to a modest increase in the overall expansion of the peptide, apparently due to charge-charge repulsion. Therefore, the conformational properties of the peptide itself are unlikely a significant contributor to the reduced LLPS propensity. Instead, analysis of the residue-residue contact probability map suggests that aromatic residues are likely the main driver of LLPS. As shown in Figure 4C for WT GY-23, the most dominant intermolecular contacts in the condensate exclusively involve histidine (H), phenylalanine (F), and tyrosine (Y) residues. H-to-K mutations do not only remove favorable *π*-*π* stacking interactions but also introduce extra charge-charge repulsion forces. It is conceivable that replacing any of these aromatic residues could significantly reduce condensate stability. Examination of the contact probability maps of condensates of the single H-to-K mutants at 280 K, where phase separation can still occur, reveals a significant reduction in the probabilities of intermolecular contacts involving the mutation site (Figure S9). Intriguingly, it appears that the location of the H-to-K mutation may have differential impacts on the condensate. Mutations at the middle region (H12K and H15K) appear to be less disruptive compared to those near the termini (H2K and H20K). Indeed, H12K and H15K can be more readily driven to phase separation, by increasing salt concentration and/or increasing pH, compared to H2K and H20K.^108^ The difference in sensitivity H-to-K mutation at central vs terminal positions is likely an entropic one, that disrupting the terminal intermolecular contacts may more significantly increase the peptide flexibility and hinder the formation of intermolecular contracts. Taken together, the apparent ability of HyRes-GPU to capture these nontrivial effects of GY-23 mutations further supports that the model is well-balanced among structural propensities and various polar and nonpolar interactions.

### Phase separation of TDP-43: correlation between helicity and condensate stability

RNA-binding protein TDP-43 is a key component of ribonucleoprotein granules and has been linked to diseases such as amyotrophic lateral sclerosis (ALS), frontotemporal dementia, and Alzheimer’s.^115–117^ Particularly, a conserved region (CR, residues 311-361) within the disordered C-terminal domain (CTD) of TDP-43 is crucial for the function of TDP-43 and harbors several ALS-associated mutations.^118^ TDP-43 CR is partially helical and can independently drive LLPS.^119,120^ Importantly, recent NMR studies further revealed that the level of residual helicity of various CR mutations, including ALS-associated ones, is positively correlated with the stability of the condensate.^75, 84^ It has been postulated that transient helices may play a direct role in mediating intermolecular interactions during phase separation.^121–123^ However, existing CG simulations using Cα-only models can not directly describe backbone-mediated interactions and residual helices during LLPS, and atomistic simulations cannot adequately sample the dynamic phase equilibrium to derive reliable intermolecular interactions within the condensate or predict the condensate stability.^75^ As such, the molecular details that underlie the apparent correlation between residual helicity and condensate stability remain poorly understood.

We first examine the ability of HyRes-GPU to recapitulate the effects of mutations on the residual helicity of four selected mutations. The results show that the residue helicity profiles predicted by HyRes-GPU are highly consistent with secondary NMR chemical shifts for the WT peptide and all four variants (Figure 6A). In particular, for WT TDP-43 CR, the simulation correctly predicts a major partial helix in residues 321-334, with residue helicity max ∼40%, and a second minor partial helix in residues 338-346 with a lower helicity of ∼20%. This is in quantitative agreement with the NMR secondary chemical shift analysis.^75^ Mutations G335A and G338A extend the major and minor partial helices, respectively, which is again in excellent agreement with NMR secondary chemical shifts. Mutation A326P reduces the helicity of the major partial helix, while M337P, located in a coiled region, has no impact on the residue helicity profile. We note that no *ad hoc* tuning is required in the above simulations. HyRes-GPU is a transferable CG protein force field that can at least semi-quantitatively describe the sequence-dependent secondary structure propensities, as well as the over peptide chain dimension (e.g., Figure 2).

**Figure 6.**
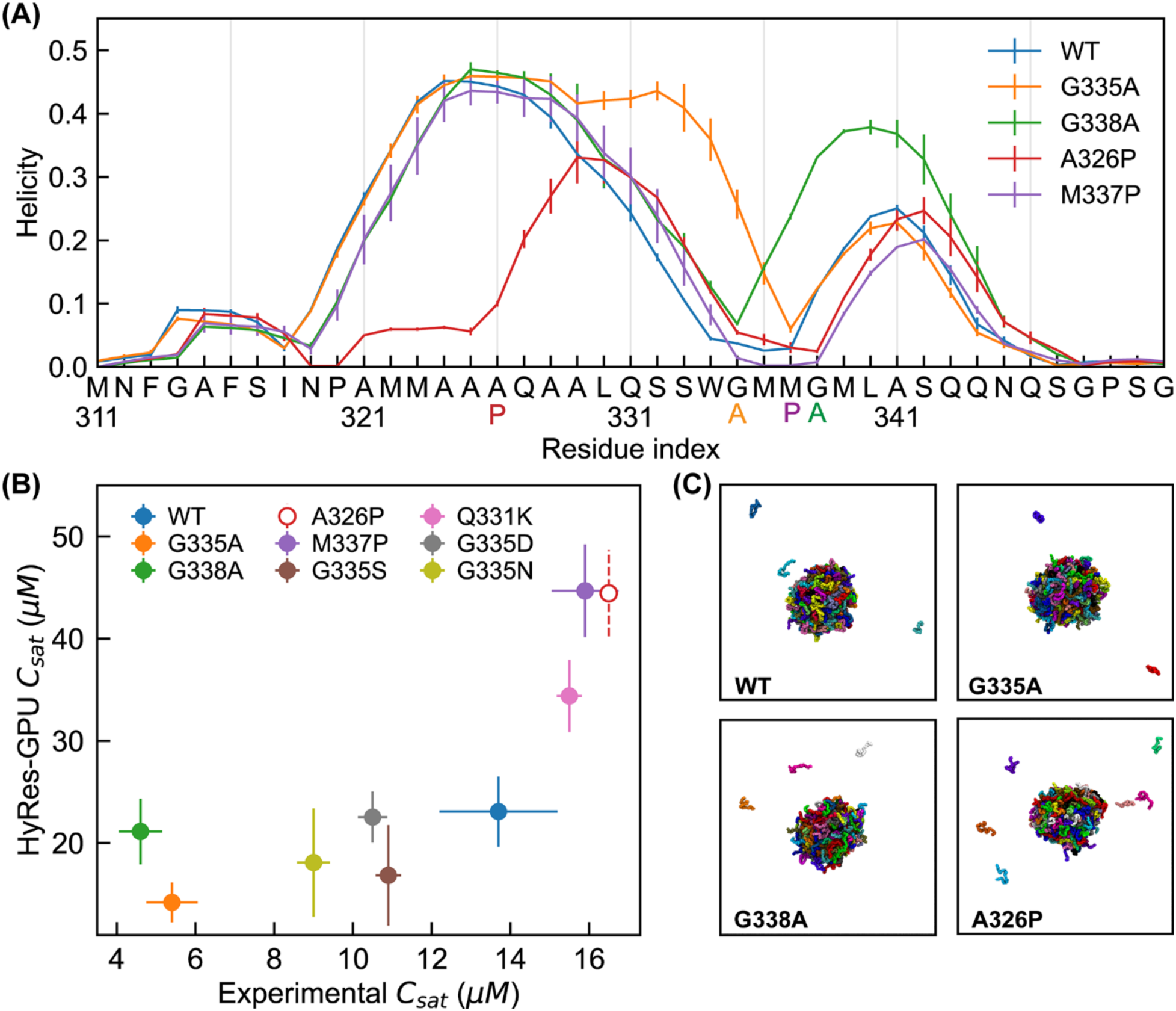
Residual helicity and LLPS propensity of TDP-43 CR variants. (A) Residue helicity profiles of WT TDP-43 CR and mutants G335A, G338A, M337P, and A326P. The error bars were calculated as the standard deviation of block averages. (B) Comparison between experimental and simulated *C*_sat_ values. The experimental values were taken from Ref. ^75^. Quantitative experimental value was not reported for mutant A326P but was stated to be similar to that of M337P. The simulated values were derived from the last 500 ns of HyRes-GPU trajectories at 300 K, and the error bars are the standard deviation calculated from three replicas. The Pearson’s correlation coefficient is *r* = 0.77 (0.85 without including G338A). (C) Representative snapshots of phase co-existence of WT, G335A, G338A, and A326P TDP-43 CR at 300 K.

We then directly simulated the phase equilibrium of WT TDP-43 CR and several single mutations using HyRes-GPU, starting from preformed high-density configurations (see Methods and Figure S2). Movies of representative simulation trajectories of WT, G335A, and A326P systems are provided in the Supplementary Materials (Movies S3, S4, and S5). Despite the substantially larger size, HyRes-GPU allows relatively fast exchange of peptides between the dilute and condensed phases. The rapid exchange is critical for accurate quantification of *C*_sat_ because of the high stability of TDP-43 CR condensates. For example, experimental *C*_sat_ is ∼15 *µ*M for the WT peptide, which translates to only ∼2 peptides in the dilute phase in the 60 x 60 x 60 nm^3^ simulation box. Figure 6B compares *C*_sat_ values from HyRes-GPU simulations and experiments for WT TDP-43 CR as well as 8 mutants, with numerical values provided in Table S4. Representative snapshots of the phase co-existence are given in Figure 6C. Overall, there is a strong correlation between the simulation and experiment (*r* = 0.77 for all and 0.85 without including G338A), even though HyRes-GPU predicts *C*_sat_ about 2-3-fold of the experimental values. The modest systematic over- estimation of *C*_sat_ could reflect limitations of HyRes-GPU, but may also be attributed to the longer protein construct characterized in the experiment study (the whole CTD instead of CR alone).^75^ Importantly, the simulations successfully recapitulate the apparent positive correlation between residual helicity and condensate stability of TDP-43. Disruption of partial helices through A326P mutation significantly increases *C*_sat_, while enhancing partial helices via G335A (or G338A) reduces *C*_sat_. This is noteworthy given the transferable and CG nature of HyRes-GPU.

### How residual helicity modulates TDP-43 CR LLPS propensity

The ability of HyRes-GPU to correctly reproduce the apparent correlation between residual helicity and *C*_sat_ allows us to further analyze the underlying molecular basis. Examination of the probability map of residue-residue contacts in the condensate of WT TDP-43 CR, shown in Figure 7A, reveals that three aromatic residues, F313, F316, and W334, play dominant roles in mediating intermolecular interactions. Previous in vitro and in vivo experiments also suggest that aromatic residues play critical role in driving the phase separation of TDP-43.^124–127^ It is noteworthy that the helical regions do not appear to be a major contributor to intermolecular contacts in the condensed phase. Inspection of the distribution of partial helices within the condensates reveals that helical regions spread around uniformly in the condensed phase and there is no apparent tendency for preferential helix-helix contacts (e.g., see Figure S10A). To quantify the fraction of residues involved in intermolecular contact when both belong to partial helices, we plot the fraction of helix-helix contacts for each residue of WT, G335A, G338A, A326P, and M337P in Figure S10B. Overall, the helix-helix contacts made only small contributions to the phase separation, up to 6% for residues in the major partial helical region for G335A. Importantly, the fractions of helix-helix contacts are strictly proportional to the residue helicity (Figure S10C). There is no apparent cooperativity in helix-helix contacts. That is, increasing helicity does not directly enhance intermolecular contact formation to help drive phase transition. This notion is actually consistent with the observation that M337P, which does not change helicity, has a similar effect on LLPS compared to A326P.

**Figure 7.**
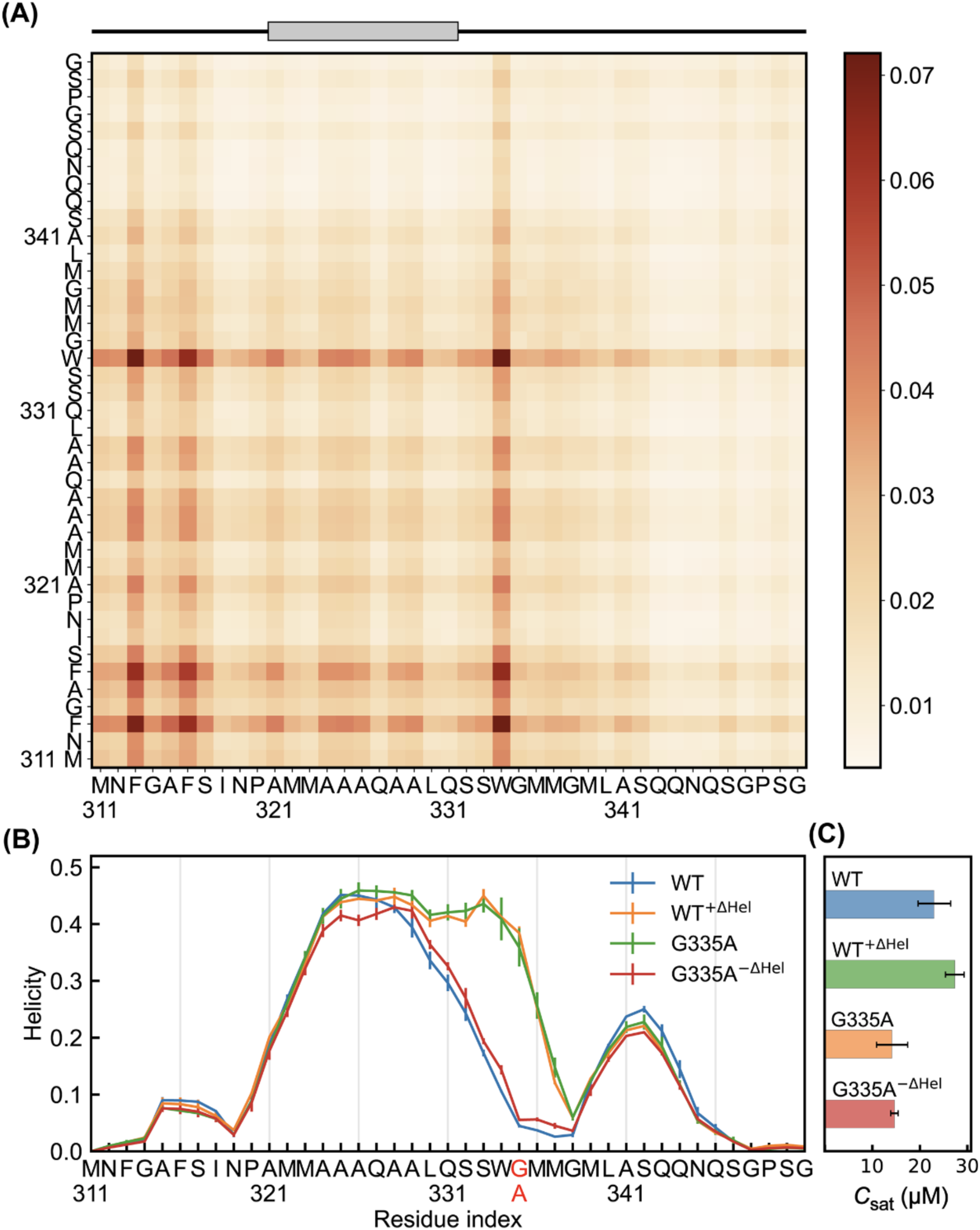
Contact map and relationship between helicity and LLPS behavior of TDP-43 CR. (A) Probabilities of intermolecular residue-residue contacts in the condensed phase of WT TDP-43 CR at 300 K, derived from the last 500 ns of the trajectories. The major partial helix is shown on top. (B) Single-chain helicity profiles of WT, WT^+ΔHel^, G335A, and G335A^-ΔHel^. (C) *C*sat of WT, WT^+ΔHel^, G335A, and G335A^-ΔHel^ from the last 500 ns of the co-existence simulations at 300 K. All error bars were calculated as the standard deviations of three replicas.

Besides affecting residual structures, mutations have direct impacts on intermolecular interactions by introducing different side chain groups. For example, the G-to-A mutation adds a hydrophobic methyl group. In addition, changing the helicity has a direct impact on the availability of backbone for intermolecular interactions. These two factors can compete with one another to influence the phase separation in a nontrivial way. To evaluate how side chain hydrophobicity and backbone availability contribute to TDP-43 CR LLPS, we designed two artificial mutants named as WT^+ΔHel^ and G335A^-ΔHel^ (Figure 7B). Here, additional dihedral restraints were imposed at residue 335 to artificially increase the helicity of WT to the same level of G335A (WT^+ΔHel^), or to reduce the helicity of G335A to the level of WT (G335A^-ΔHel^). The simulations show that WT^+ΔHel^ actually reduces the condensate stability despite the increased helicity, resulting in an increased *C*_sat_ (Figure 7C). This is opposite of what one may expect if helicity has a direct and positive contribution to condensate stability. Instead, artificially increasing helicity in the WT sequence reduces the availability of backbone for intermolecular interactions, destabilizing the condensate. This is supported by analysis of residue-residue contact probabilities, showing backbone-backbone contacts are the most affected in WT^+ΔHel^ (Figure S11). For G335A^-ΔHel^, the calculated *C*_sat_ is similar to that of G335A despite reduced helicity near residue 335 (Figure 7C). The implication is that introduction of the hydrophobic Ala side chain is the dominant contributor behind increased condensate stability. This effect becomes evident when comparing contact probabilities between WT and G335A^-ΔHel^ (Figure S12A), as well as between WT^+ΔHel^ and G335A (Figure S12B). Remarkably, the hydrophobic Ala side chain substantially augments contacts of residues near position 335, even with different levels of residual helicities.

For A326P and M337P, the proline mutation leads to a global reduction in the contact probabilities (e.g., see Figure S13), presumably because prolines reduce the ability of peptides to pack closely in the condensed phase. As such, M337P can lead to a similar increase in *C*_sat_, even though its helicity is very similar to that of WT. Taken together, these analyses strongly suggest that mutations will perturb the complex balance among multiple factors, including peptide conformation, backbone availability, and sidechain hydrophobicity, to modulate the IDP phase equilibrium. The nontrivial net effects of mutations on the balance are likely responsible for the contrasting observations regarding the influence of helicity on the propensity for liquid-liquid phase separation (LLPS) in various experimental studies.^75, 128^ HyRes-GPU should provide a powerful tool for studying these nontrivial effects with the ability to accurately describe residual structures and backbone-mediated interactions.

## Conclusion

In summary, we have developed an efficient GPU-accelerated HyRes CG protein model that provides atomistic description of the peptide backbone to enable more accurate simulations of phase separation of IDPs. The model is physics-motivated and transferable. It provides a semi-quantitative description of both local residual structures and overall level of compaction of the monomeric form of a diverse set of IDPs. Application to two phase-separating IDPs, GY-23 and TDP-43 CR, demonstrates that HyRes-GPU is sufficiently balanced to capture rapid peptide exchange between the dilute and condensed phases and recapitulate the effects of single mutations on the dynamic phase equilibrium. Our in-depth analysis of GY-23 and TDP-43 CR phase separation also provides new insights on the role of backbone-mediated interactions and coupling of residual structures with phase separation. In the case of highly disordered GY-23, it is found that phase separation can promote the formation of *β*-structures within the condensed phase to augment intermolecular interactions and stabilize the condensates. For TDP-43 with substantial residual helicity, HyRes-GPU simulations reveal that phase separation is mainly driven by aromatic residues and there are minimal preferential helix-helix interactions in the condensate. Instead, the apparent positive correlation between helicity and condensate stability, observed in both experiment and simulations, is a net result of nontrivial effects of perturbing peptide conformations, backbone availability, and sidechain hydrophobicity. In particular, increasing helicity itself actually reduces the availability of backbone for intermolecular interactions and can destabilize the condensate. Taken together, HyRes-GPU dramatically expands the capability of CG simulations of IDP phase separation and can be deployed to elucidate how detailed structural and sequence features may affect the phase equilibrium at the molecular level.

## Methods

### HyRes simulations of IDP monomers

A total of 26 IDPs ranging from ∼20 to ∼300 residues were simulated in this work. The sequences of these IDPs are provided in Table S1, with experimental radii of gyration and solution conditions summarized in Table S2. The N- and C-termini of all proteins are capped with acetyl and N-methyl amide groups, respectively. All simulations were carried out using CHARMM^129^ and OpenMM^130^ on GPUs. The initial extended structures were generated using CHARMM and then energy was minimized. Langevin dynamics was performed with a collision frequency of 0.2 ps^-1^ and an MD timestep of 4 fs. The salt concentration is set at 150 mM, consistent with most experimental conditions (Table S3). All bonds involving hydrogen atoms were constrained by the SHAKE algorithm. Nonbonded interactions were smoothly switched off from 1.6 nm to 1.8 nm. All monomers were simulated for 1 *µs* at 300 K in HyRes unless otherwise noted. Additional simulations of GY-23 mutants were simulated for 400 ns per replica, which proved to be long enough for achieving sufficient convergence (e.g., see error bars in Figure S7). Four additional 400-ns production simulations were also performed for WT GY-23 monomer at 260 K, 270 K, 280 K, and 290 K to examine the temperature dependence of conformational properties in comparison to results from condensate simulations.

The first 200 ns trajectories were excluded in subsequent structural analysis, which was performed using a combination of CHARMM and in-house scripts. The secondary structures were identified using the standard Dictionary of Secondary Structure of Proteins (DSSP) protocol within the MDTraj package.^131^ *R*_g_ values from HyRes-GPU simulations are uniformly shifted up by 4 Å to roughly account for larger side chain beads as done previously. Viscosity of GY-23 peptides in dilute and condensed phases was calculated from MSD correlation functions of the C_α_ atom of the central residue (residue 12). The error bars for all properties were estimated from differences between results calculated from two independent simulation replicas. All molecular visualizations were done using VMD.^132^

### Simulation of phase separation and equilibrium

To prepare the initial configuration for LLPS simulation of GY-23, we first generated a minimized extended conformation and then packed 200 copies together using the Packmol Software.^133^ The initial cubic box was used for packing 30*30*30 nm^3^ with a distance tolerance of 10 Å to avoid atom overlap (Figure S2A). The initial packed state was subject to 1-ns constant pressure and temperature (NPT) equilibration at 300 K, which would result in a high-density compact configuration of ∼12 nm in diameter. The cubic box size was subsequently increased to 45 nm for production simulation of phase equilibrium. Multiple copies of constant volume and temperature (NVT) simulations were then performed at every 5 K from 260 K to 320 K for 2 *µs*, which have been shown to be sufficient to reach phase equilibrium (see main text). A separate NVT simulation was performed at 600 K to generate a fully dispersed initial configuration (Figure 2A), which was then used to initiate three independent 2-*µs* simulations of spontaneous phase separation of WT GY-23 at 300 K. The convergence of the phase equilibrium will be examined by comparing simulations initiated from compact and dispersed initial configurations (see main text). Additional simulations were performed for WT GY- 23 to examine the finite-size efforts on the phase equilibrium, using two additional constructs with 100 copies in a 35.2-nm cubic box and 300 copies in a 51.7-nm cubic box (Figure S4). Similar protocols were used to generate the initial compact configurations and the simulations were performed for 2 -*µ*s at 300 K. A similar protocol was used to generate the compact initial configuration of TDP-43 CR (Figure S2B), except for the larger packing box used (45*45*45 nm^3^). The phase equilibrium simulations of TDP-43 CR were performed at 300 K in a 60-nm cubic box and lasted 2 *µs* each.

### Phase diagram and properties of the condensates

A single dominant droplet was observed in all phase equilibrium simulations. They were first identified using the density-based spatial clustering of applications with noise (DBSCAN) clustering algorithm within the Python Scikit package.^134^ Density profiles show that the density of the condensate is uniform within the droplet (e.g., Figure 3D). We chose a cutoff radius of 3 nm to determine the concentration of the condensates of GY-23, averaged over the last 500 ns were used, where the phase equilibrium is fully established (e.g., see Figure S3). Residue-residue contact maps were determined based on the minimum distance of two residues (only heavy atoms), where the contact was identified if the minimum distance was no longer than 0.5 nm. The critical points, *T*_c_ and, *ρ*_c_ were determined by fitting the calculated densities at different temperatures to the following Ising-like model,^64, 114^

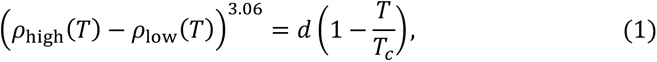

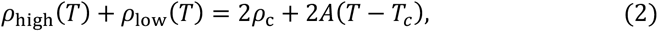

where *ρ*_high_ and *ρ*_low_ are the densities of condensed and dilute phases. The critical exponent is set to be 3.06 for the universality class of 3D Ising model. *d* and *A* are protein-specific fitting parameters.

## Supporting information

All SI Tables and Figures

## Acknowledgments

We thank Peter Eastman for his generous help with the modification of OpenMM to provide a more efficient implementation of the hydrogen-bonding term. This modification has been merged into the latest version of OpenMM, available at github.com/openmm/openmm. All simulations were performed on the Pikes GPU cluster housed in the Massachusetts Green High-Performance Computing Cluster (MGHPCC). This work was supported by the National Institutes of Health Grant R35 GM144045 (J. C.).

## Author contributions

Y. Z., S. L. and J. C., conception and design of the study; Y. Z., S.L., performing the simulation and analysis; X. G. and S. L. implementation and performance analysis of HyRes on GPU; Y. Z., S. L. and J. C., analysis and interpretation of data, drafting and revising the manuscript.

**Supporting Information** accompanies this paper at doi: xxxx.

