## Supplementary material for "Accurate Simulation of Coupling between Protein Secondary Structure and Liquid-Liquid Phase Separation": All SI Tables and Figures

### Supplementary Tables

**Table S1.** Lennard-Jones parameters for all backbone atoms in HyRes-GPU.

| Atom Type | $\varepsilon_i$ (kcal/mol) | $\varepsilon_i$ (1-4) (kcal/mol) | $R_{min}$ (Å) |
|---|---|---|---|
| **H** | - 0.0149 | - 0.0498 | 0.225 |
| **C** | - 0.0180 | - 0.1000 | 2.100 |
| **Cα** | - 0.0073 | - 0.1000 | 2.365 |
| **Cα (Gly)** | - 0.0172 | - 0.1000 | 2.235 |
| **N** | - 0.0356 | - 0.2384 | 1.600 |
| **O** | - 0.2390 | - 0.1159 | 1.600 |

**Table S2.** Sequences of all IDPs studied in the work. Peptides that have already been shown to drive phase separation are colored in blue.

| Protein | Sequence |
|---|---|
| **(AAQAA)$_3$** | AAQAA AAQAA AAQAA |
| **KID** | TDSQK RREIL SRRPS YRKIL NDLSS DAP |
| **α-synuclein** | MDVFM KGLSK AKEGV VAAAE KTKQG VAEAA GKTKE GVLYV GSKTK<br>EGVVH GVATV AEKTK EQVTN VGGAV VTGVT AVAQK TVEGA GSIAA<br>ATGFV KKDQL GKNEE GAPQE GILED MPVDP DNEAY EMPSE EGYQD<br>YEPEA |
| **ACTR** | GTQNR PLLRN SLDDL VGPPS NLEGQ SDERA LLDQL HTLLS NTDAT<br>GLEEI DRALG IPELV NQGQA LEPKQ D |
| **Ash1** | GASAS SSPSP STPTK SGKMR SRSSS PVRPK AYTPS PRSPN YHRFA<br>LDSPP QSPRR SSNSS ITKKG SRRSS GSSPT RHTTR VCV |
| **hNHE1cdt** | MVPAH KLDSP TMSRA RIGSD PLAYE PKEDL PVITI DPASP QSPES<br>VDLVN EELKG KVLGL SRDPA KVAEE DEDDD GGIMM RSKET SSPGT<br>DDVFT PAPSD SPSSQ RIQRC LSDPG PHPEP GEGEP FFPKG Q |
| **IBB** | GCTNE NANTP AARLH RFKNK GKDST EMRRR RIEVN VELRK AKKDD<br>QMLKR RNVSS FPDDA TSPLQ ENRNN QGTVN WSVDD IVKGI NSSNV<br>ENQLQ AT |
| **K18** | MQTAP VPMPD LKNVK SKIGS TENLK HQPGG GKVQI INKKL DLSNV<br>QSKCG SKDNI KHVPG GGSVQ IVYKP VDLSK VTSKC GSLGN IHHKP<br>GGGQV EVKSE KLDFK DRVQS KIGSL DNITH VPGGG NKKIE |
| **N49** | GCQTS RGLFG NNNTN NINNS SSGMN NASAG LFGSK P |
| **N98** | GCFNK SFGTP FGGGT GGFGT TSTFG QNTGF GTTSG GAFGT SAFGS<br>SNNTG GLFGN SQTKP GGLFG TSSFS QPATS TSTGF GFGTS TGTAN<br>TLFGT ASTGT SLFSS QNNAF AQNKP TGFGN FGTST SSGGL FGTTN<br>TTSNP FGSTS GSLFG P |
| **NLS** | ACETN KRKRE QISTD NEAKM QIQEE KSPKK KRKKR SSKAN KPPE |
| **NSP** | GCNFN TPQQN KTPFS FGTAN NNSNT TNQNS STGAG AFGTG QSTFG<br>FNNSA PNNTN NANSS ITPAF GSNNT GNTAF GNSNP TSNVF GSNNS<br>TTNTF GSNSA GTSLF GSSSA QQTKS NGTAG GNTFG SSSLF NNSTN<br>SNTTK PAFGG LNFGG GNNTT PSSTG NANTS NNLFG ATANA N |
| **NUL** | GCGFK GFDTS SSSSN SAASS SFKFG VSSSS SGPSQ TLTST GNFKF<br>GDQGG FKIGV SSDSG SINPM SEGFK FSKPI GDFKF GVSSE SKPEE<br>VKKDS KNDNF KFGLS SGLSN PV |
| **NUS** | GCPSA SPAFG ANQTP TFGQS QGASQ PNPPG FGSIS SSTAL FPTGS<br>QPAPP TFGTV SSSSQ PPVFG QQPSQ SAFGS GTTPN |
| **p53 (1-93)** | MEEPQ SDPSV EPPLS QETFS DLWKL LPENN VLSPL PSQAM DDLML<br>SPDDI EQWFT EDPGP DEAPR MPEAA PPVAP APAAP TPAAP APAPS<br>WPL |

| | |
|---|---|
| **ProTα** | MSDAA VDTSS EITTK DLKEK KEVVE EAENG RDAPA NGNAE NEENG EQEAD NEVDE EEEEG GEEEE EEEEG DGEEE DGDED EEAES ATGKR AAEDD EDDDV DTKKQ KTDED D |
| **SH4-UD** | MGSNK SKPKD ASQRR RSLEP AENVH GAGGG AFPAS QTPSK PASAD GHRGP SAAFA PAAAE PKLFG GFNSS DTVTS PQRAG PLAGG |
| **Sic1** | GSMTP STPPR SRGTR YLAQP SGNTS SSALM QGQKT PQKPS QNLVP VTPST TKSFK NAPLL APPNS NMGMT SPFNG LTSPQ RSPFP KSSVK RT |
| **Hst5** | DSHAK RHHGY KRKFH EKHHS HRGY |
| **$(Hst5)_2$** | DSHAK RHHGY KRKFH EKHHS HRGYD SHAKR HHGYK RKFHE KHHSH RGY |
| **FUS** | MASND YTQQA TQSYG AYPTQ PGQGY SQQSS QPYGQ QSYSG YSQST DTSGY GQSSY SSYGQ SQNTG YGTQS TPQGY GSTGG YGSSQ SSQSS YGQQS SYPGY GQQPA PSSTS GSYGS SSQSS SYGQP QSGSY SQQPS YGGQQ QSYGQ QQSYN PPQGY GQQNQ YNS |
| **LAF-1 RGG (WT)** | MESNQ SNNGG SGNAA LNRGG RYVPP HLRGG DGGAA AAASA GGDDR RGGAG GGGYR RGGGN SGGGG GGGYD RGYND NRDDR DNRGG SGGYG RDRNY EDRGY NGGGG GGGNR GYNNN RGGGG GGYNR QDRGD GGSSN FSRGG YNNRD EGSDN RGSGR SYNND RRDNG GDG |
| **LAF-1RGG (Shuffled)** | MNNSG DNDRG SGNYG LRNSF GDDGY GDNGN DEGNS GYRNR GLGGD RADEY GNSGG NGDNE AAPNA SDRDD AHYYD SDDYD DGGGG RGSGG AGGGG ARGPG SNRAG RYGGG GRRGR GRGNG YNGNR SQRRR GGGRG RGNRG YRVGN GNGQS GGRNS RGGGG GNGGA NYGLE HHHHH H |
| **A1 LCD** | GSMAS ASSSQ RGRSG SGNFG GGRGG GFGGN DNFGR GGNFS GRGGF GGSRG GGGYG GSGDG YNGFG NDGSN FGGGG SYNDF GNYNN QSSNF GPMKG GNFGG RSSGG SGGGG QYFAK PRNQG GYGGS SSSSS YGSGR RF |
| **TDP-43 CTD** | RQLER SGRFG GNPGG FGNQG GFGNS RGGGA GLGNN QGSNM GGGMN FGAFS INPAM MAAAQ AALQS SWGMM GMLAS QQNQS GPSGN NQNQG NMQRE PNQAF GSGNN SYSGS NSGAA IGWGS ASNAG SGSGF NGGFG SSMDS KSSGW GM |
| **Ddx4 LCD** | MGDED WEAEI NPHMS SYVPI FEKDR YSGEN GDNFN RTPAS SSEMD DGPSR RDHFM KSGFA SGRNF GNRDA GECNK RDNTS TMGGF GVGKS FGNRG FSNSR FEDGD SSGFW RESSN DCEDN PTRNR GFSKR GGYRD GNNSE ASGPY RRGGR GSFRG CRGGF GLGSP NNDLD PDECM QRTGG LFGSR RPVLS GTGNG DTSQS RSGSG SERGG YKGLN EEVIT GSGKN SWKSE AEGGE S |
| **GY-23** | GHGLY GAGFA GHGLH GFAGH GLY |
| **TDP-43 CR** | $M^{311}$NFGA FSINP AMMAA AQAAL QSSWG MMGML ASQQN QSGPS GNNQN $QGNMQ^{360}$ |

**Table S3.** Experimental $R_g$ and the corresponding measurement conditions including temperature ($T$), salt concentration ($C_{salt}$), and pH. The lengths of the proteins ($N_{res}$) are also listed.

| Protein | $N_{res}$ | $R_g$ (nm) | $T$ (K) | $C_{salt}$ (M) | pH |
|---|---|---|---|---|---|
| α-synuclein[1] | 140 | 3.55 | 293 | 0.2 | 7.4 |
| ACTR[2] | 71 | 2.6 | 278 | 0.2 | 7.4 |
| Ash1[3, 4] | 83 | 2.9 | 293 | 0.15 | 7.5 |
| hNHE1cdt[2] | 131 | 3.63 | 278 | 0.2 | 7.4 |
| IBB[5] | 97 | 3.20 | 277 | 0.16 | 7.4 |
| K18[5] | 130 | 3.80 | 277 | 0.16 | 7.4 |
| N49[5] | 36 | 1.59 | 277 | 0.16 | 7.4 |
| N98[5] | 151 | 2.86 | 277 | 0.16 | 7.4 |
| NLS[5] | 44 | 2.40 | 277 | 0.16 | 7.4 |
| NSP[5] | 176 | 4.10 | 277 | 0.16 | 7.4 |
| NUL[5] | 112 | 3.00 | 277 | 0.16 | 7.4 |
| NUS[5] | 115 | 2.49 | 277 | 0.16 | 7.4 |
| p53 (1-93)[6] | 93 | 2.87 | 293 | 0.15 | 6.8 |
| ProTα[7, 8] | 111 | 3.79 |  | 0.15 | 7.0 |
| SH4-UD[9] | 85 | 2.90 | 277 | 0.22 | 7.5 |
| Sic1[10] | 92 | 3.21 | 298 | 0.16 | 7.5 |
| Hst5[11] | 24 | 1.38 | 293 | 0.15 | 7.5 |
| $(Hst5)_2$[11] | 48 | 1.87 | 298 | 0.15 | 7.0 |
| FUS[12] | 163 | 3.32 | 297 | 0.15 | 7.4 |
| LAF-1 RGG (WT)[13] | 176 | 3.08 | 293 | 0.15 | 7.4 |
| LAF-1 RGG (shuffled)[13] | 176 | 3.0 | 293 | 0.15 | 7.4 |
| A1 LCD[14] | 137 | 2.76 | 298 | 0.15 | 7.0 |
| TDP-43 CTD[15] | 147 | 2.8 | 298 | 0.2 | 6.1 |
| Ddx4 LCD[16] | 236 | 3.61 | 297 | 0.15 | 6.5 |

**Table S4.** Experimental and simulated $C_{sat}$ for TDP-43 CR variants.

| TDP-43 CR | Experimental $C_{sat}$ (μM)[17] | Simulated $C_{sat}$ (μM) |
|---|---|---|
| WT | 13.7 ± 1.5 | 23.1 ± 3.4 |
| $WT^{+\Delta Hel}$ | -- | 27.4 ± 3.2 |
| G335A | 5.4 ± 0.7 | 14.2 ± 2.0 |
| $G335A^{-\Delta Hel}$ | -- | 14.7 ± 0.8 |
| G338A | 4.6 ± 0.6 | 21.1 ± 3.1 |
| A326P | -- | 44.5 ± 4.2 |
| M337P | 15.9 ± 0.9 | 44.7 ± 4.5 |
| G335S | 10.9 ± 0.3 | 16.8 ± 4.9 |
| Q331K | 15.5 ± 0.3 | 34.4 ± 3.5 |
| G335D | 10.5 ± 0.4 | 22.5 ± 2.5 |
| G335N | 9.0 ± 0.4 | 18.1 ± 5.3 |

### Supplementary Movies

**Movie S1**: Spontaneous phase separation of WT GY-23 starting from a fully dispersed initial configuration. The simulation was performed in HyRes-GPU at 300 K and lasted 2000 ns. The simulation box contains 200 copies in 45 x 45 x 45 $nm^3$, for a total concentration of ~3.6 mM.

**Movie S2**: Spontaneous phase separation of WT GY-23 starting from a preformed high-density initial configuration. The simulation was performed in HyRes-GPU at 300 K and lasted 2000 ns. The simulation box contains 200 copies in 45 x 45 x 45 $nm^3$, for a total concentration of ~3.6 mM.

**Movie S3**: Spontaneous phase separation of TDP-43 CR WT starting from a preformed high-density initial configuration. The simulation was performed in HyRes-GPU at 300 K and lasted 2000 ns. The simulation box contains 200 copies and is 60 x 60 x 60 $nm^3$, for a total concentration of ~1.5 mM.

**Movie S4**: Spontaneous phase separation of TDP-43 CR G335A starting from a preformed high-density initial configuration. The simulation was performed in HyRes-GPU at 300 K and lasted 2000 ns. The simulation box contains 200 copies and is 60 x 60 x 60 $nm^3$, for a total concentration of ~1.5 mM.

**Movie S5**: Spontaneous phase separation of TDP-43 CR A326P starting from a preformed high-density initial configuration. The simulation was performed in HyRes-GPU at 300 K and lasted 2000 ns. The simulation box contains 200 copies and is 60 x 60 x 60 $nm^3$, for a total concentration of ~1.5 mM.

### Supplementary Figures

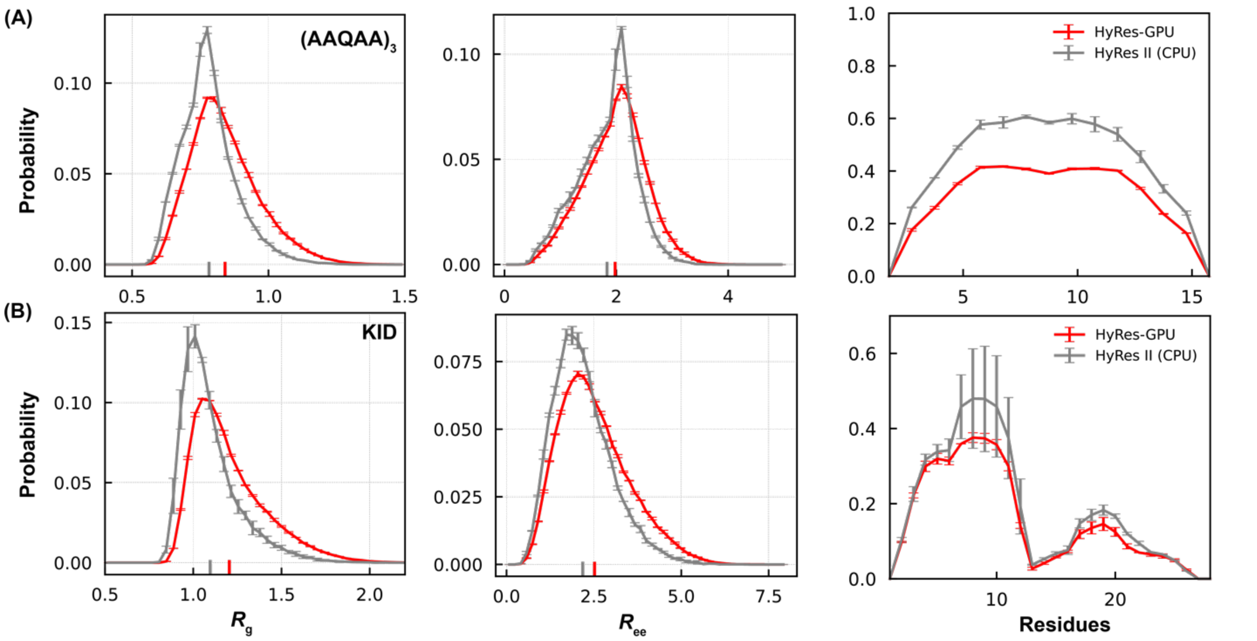


**Figure S1.** The probability distributions of $R_g$, $R_{ee,}$ and residue helicity of $(AAQAA)_3$ (A) and KID (B) sampled from HyRes-GPU and HyRes II (CPU) at 300 K.

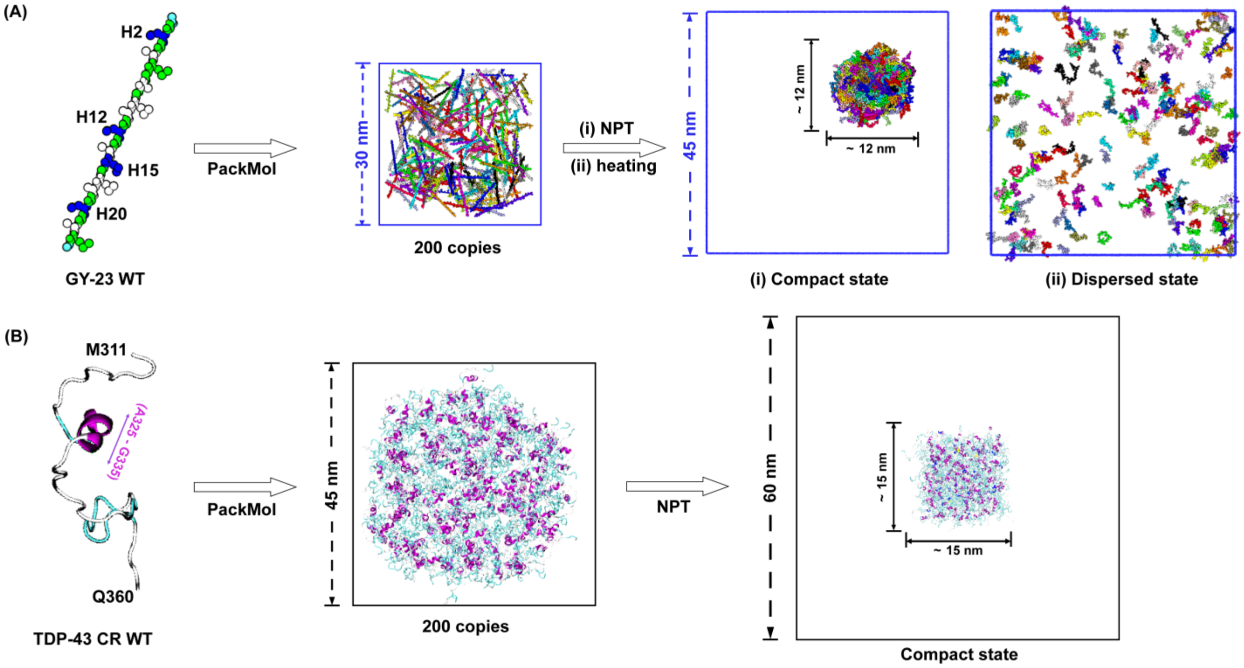


**Figure S2.** Preparation of the initial conformations for phase separation simulations of (A) GY-23 and (B) TDP-43 CR. Similar procedures were used for both WT peptides and their mutants.

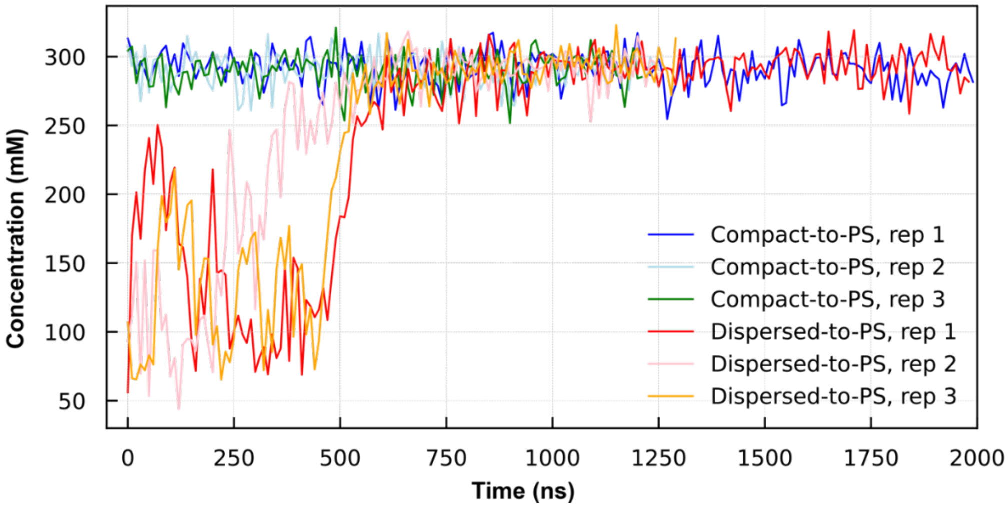


**Figure S3.** The density of the largest droplets during six simulations of WT GY-23 at 300 K initiated from the compact and dispersed initial conformation, respectively.

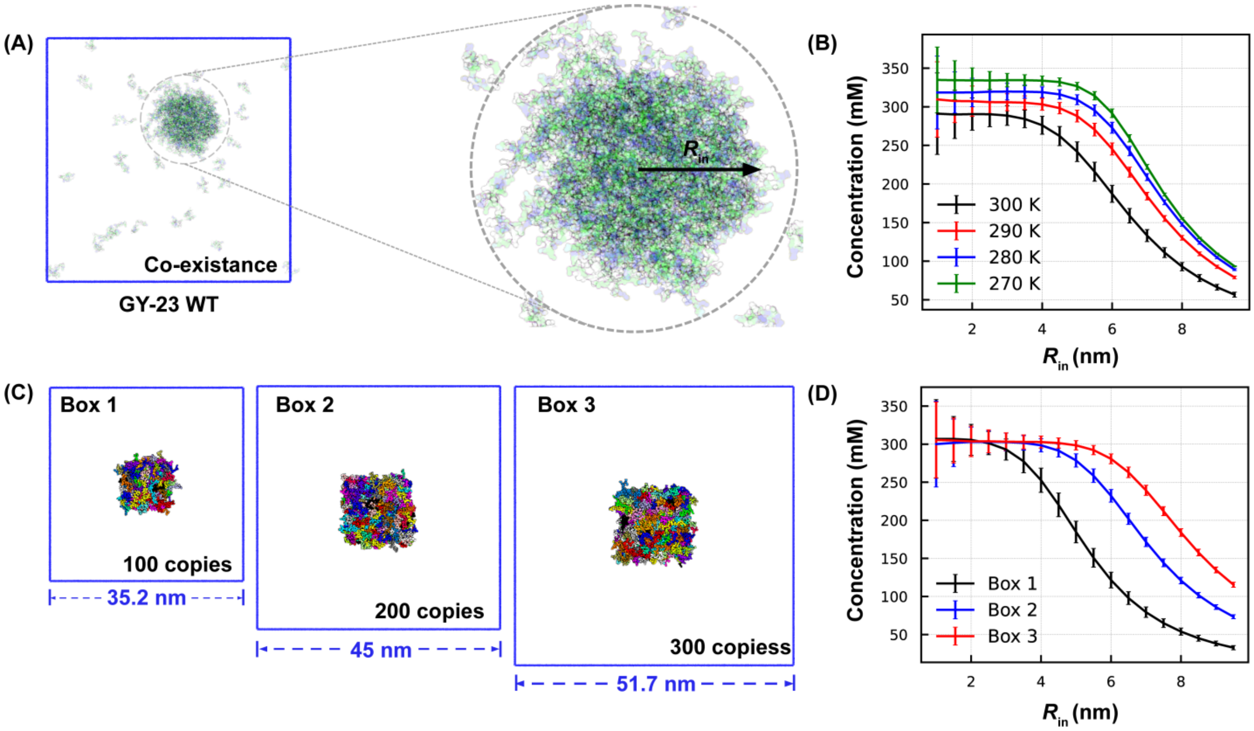


**Figure S4.** Finite-size effects on the simulation of GY-23 condensates. (A) The droplet was first identified using DBSCAN with a radius of 2.5 nm. The density profile was then calculated as a function of radius around the center of the droplet. (B) Density profiles of the final droplet of WT GY-23 at equilibrium at four selected temperatures. The box size is 45 nm. (C) The initial compact configurations WT GY-23 with three different box sizes but at the same total concentration of 3.6 mM. (D) Density profiles were calculated for droplets at phase equilibrium from simulations of WT GY-23 with three different box sizes as shown in (C).

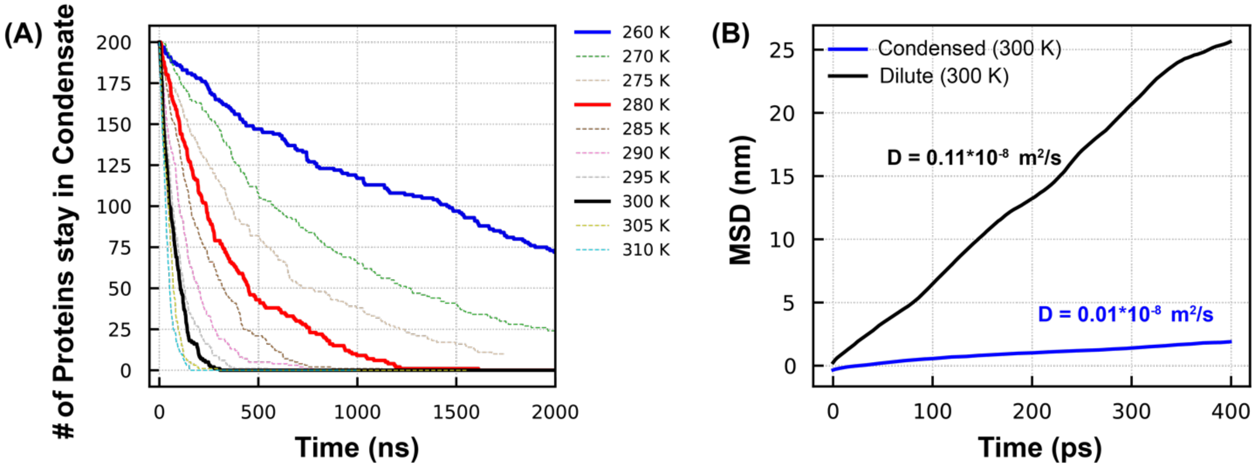


**Figure S5.** (A) Numbers of WT GY-23 peptides that remain in the preformed droplet as a function of simulation time at different simulation temperatures. (B) MSD of GY-23 in the dilute (black) and condensed (blue) phases at 300 K as a function of time.

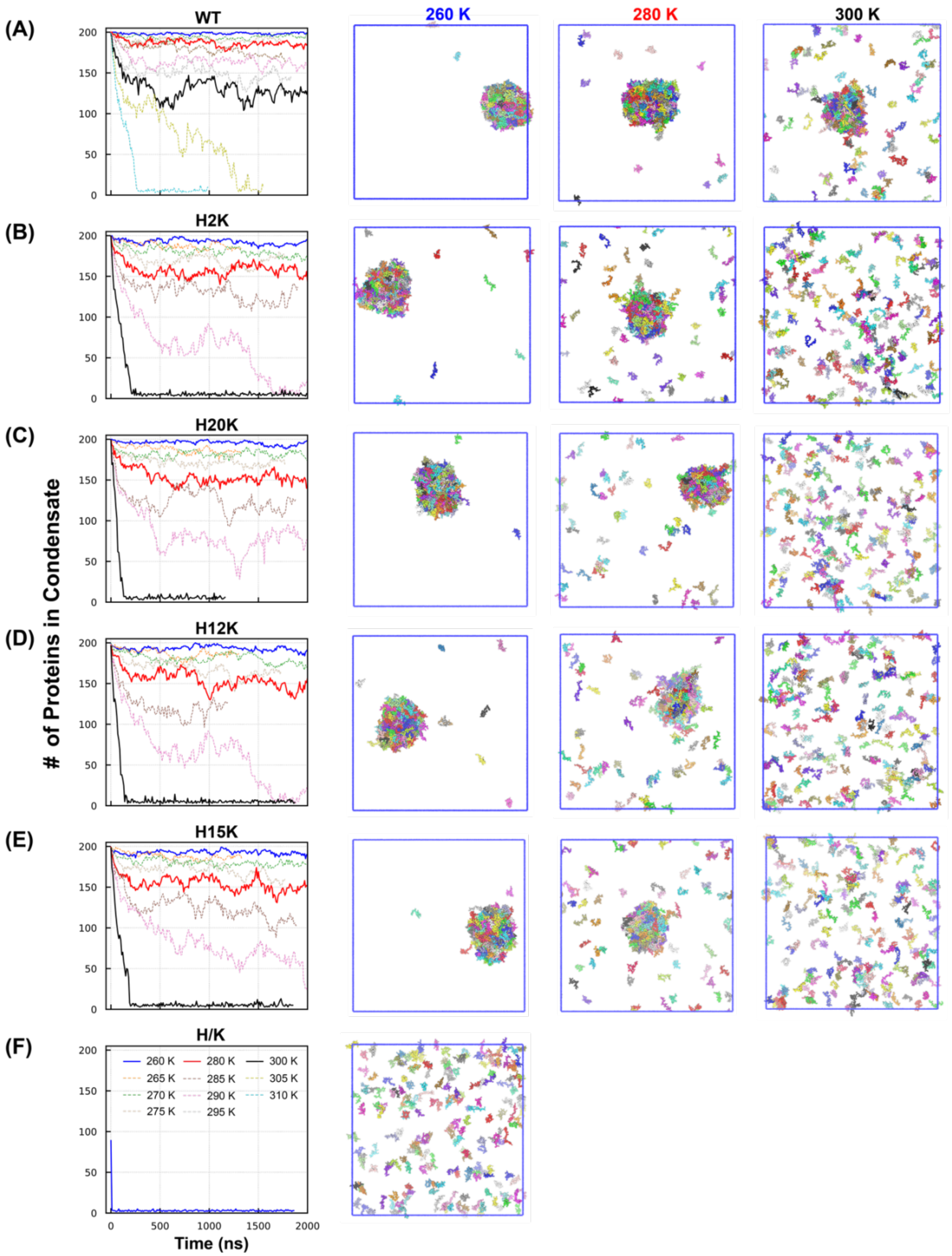


**Figure S6.** Phase separation of WT and mutant GY-23 peptides. The left column plots the number of peptides in the largest cluster as a function of simulation time at temperatures ranging from 260 up to 310 K. Traces at 260 K, 280 K and 300 K are highlighted using solid lines, with representative final snapshots shown.

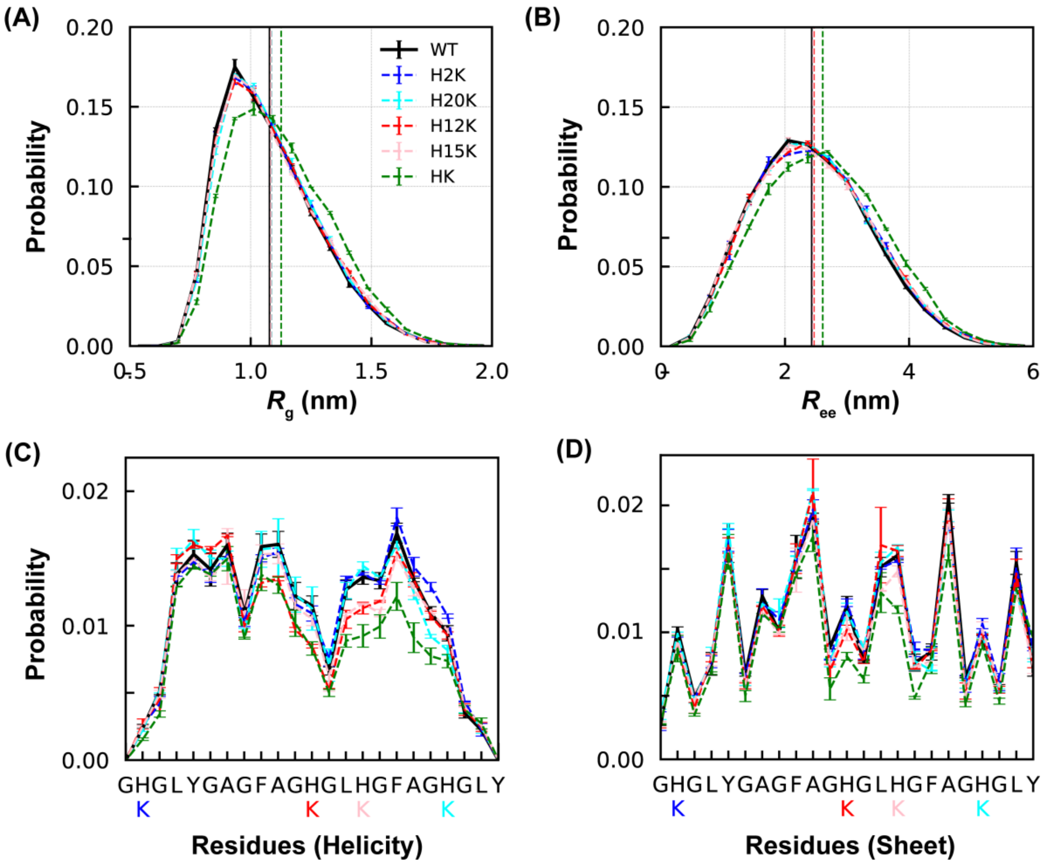


**Figure S7.** Conformational properties of monomeric GY-23 peptides. Probability distributions of (A) $R_g$, (B) $R_{ee}$, (C) average residue helicity, and (D) $\beta$ structure propensity of WT GY-23 and five mutants at 300 K. The vertical lines in panels A and B mark the average values.

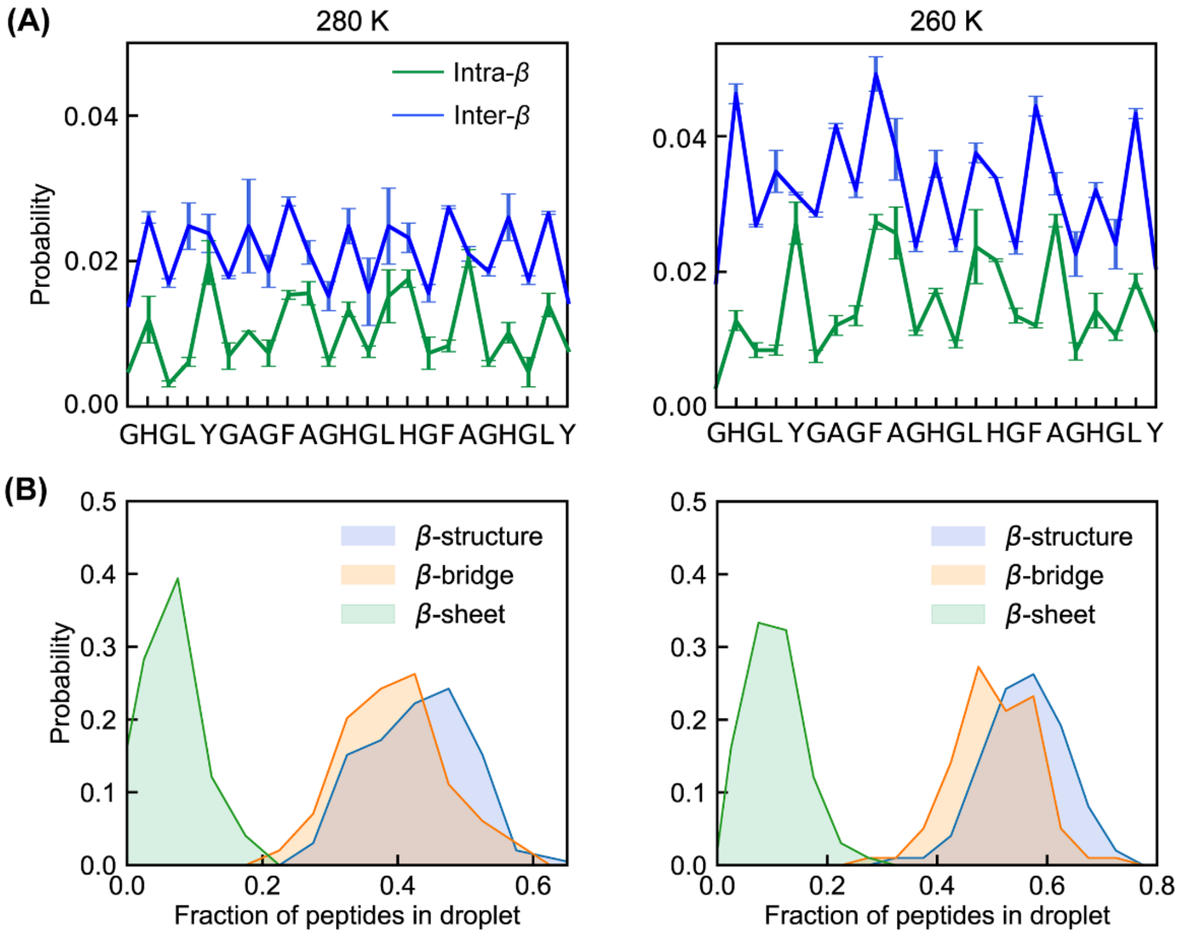


**Figure S8.** $\beta$-structure properties in the condensed phase of WT GY-23 at 280 K (left column) and 260 K (right column). (A) The propensities of intra- and inter-β structures. (B) Distributions of the fractions of peptides involved in all $\beta$-structures, $\beta$-bridges, and $\beta$-sheets in the droplet.

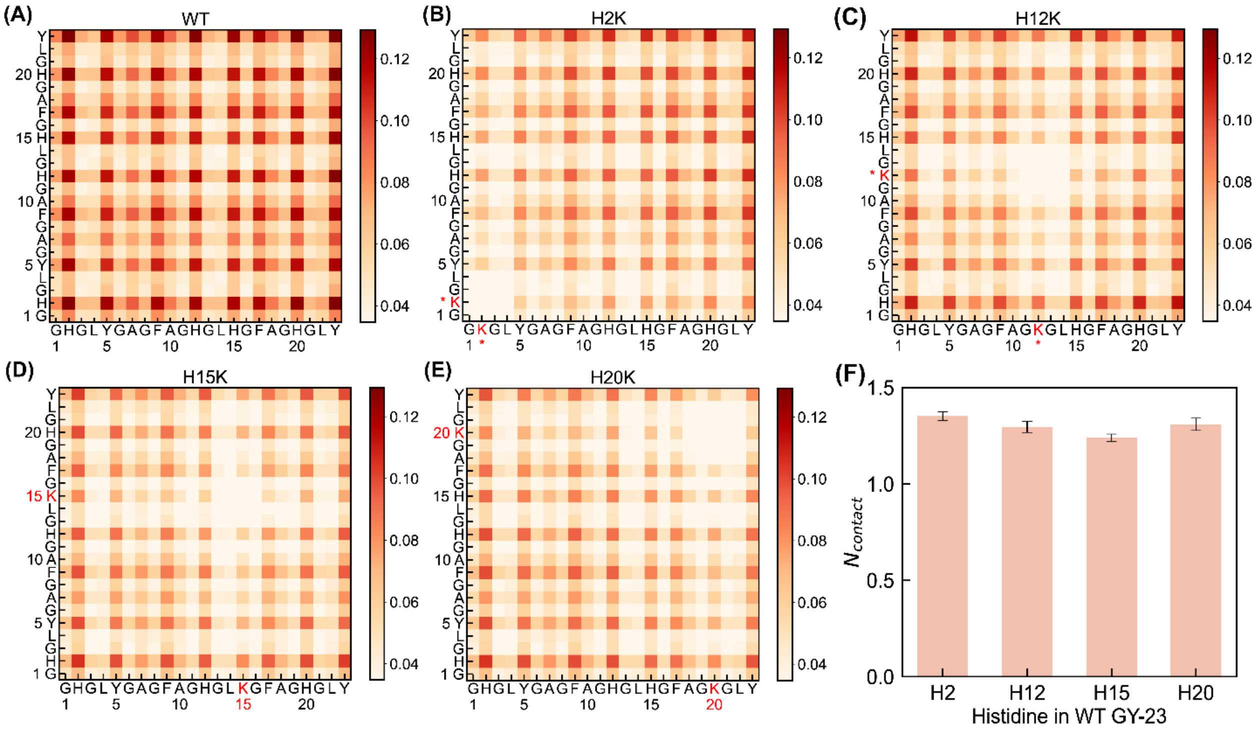


**Figure S9.** Residue contacts of GY-23 variants. Probability maps of inter-molecular residue-residue contacts in the condensed phases of (A) WT (at 300 K), and (B-E) H2K, H12K, H15K, and H20K at 280 K. (F) Average mumbers of intermolecular contacts per frame for residues H2, H12, H15, and H20 in the condensate of WT GY-23 at 300 K.

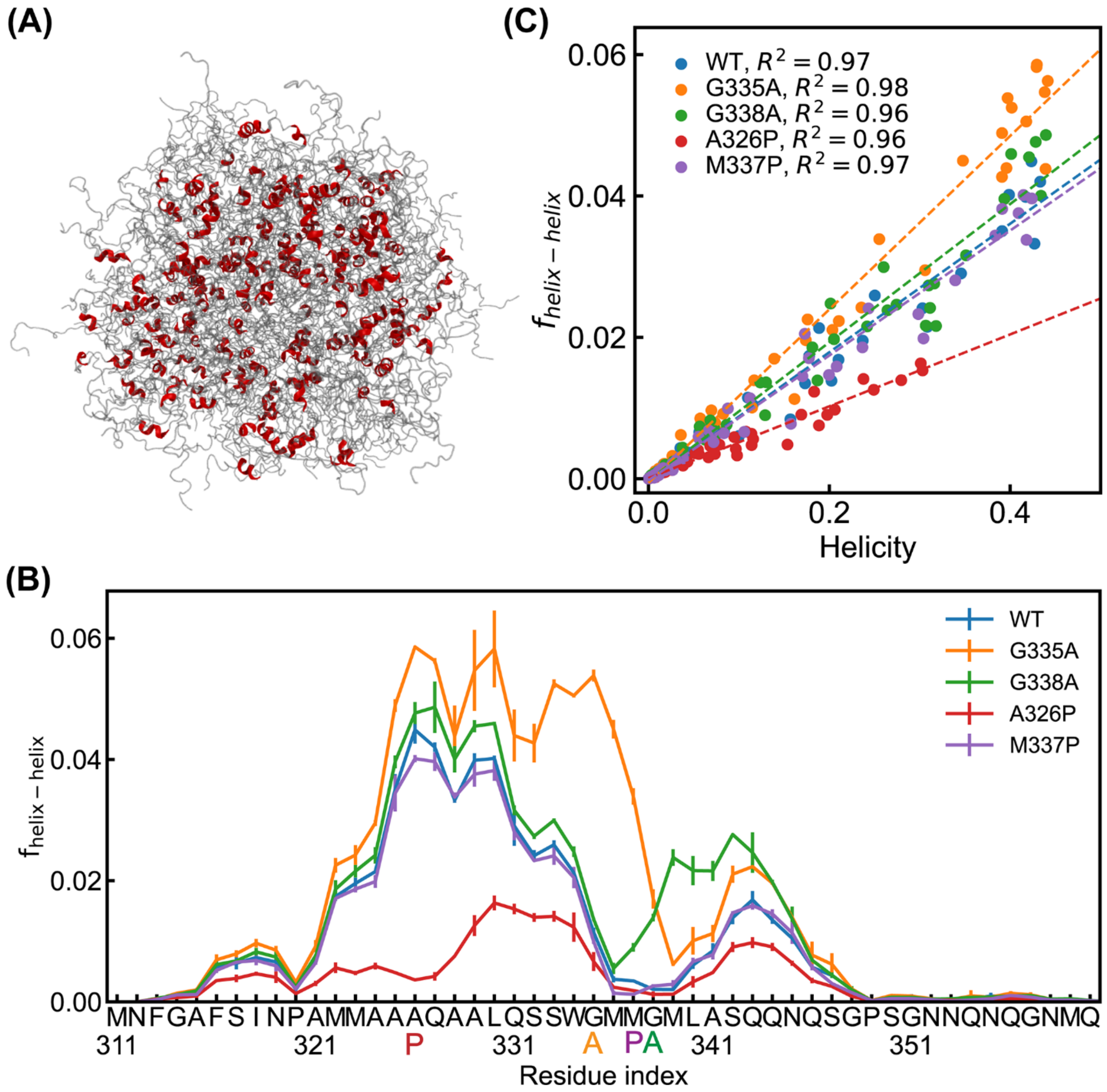


**Figure S10.** Helix-helix interactions in condensed phase. (A) A representative snapshot of the condensate of WT highlighting the presence of partial helices. Note that these partial helices are dispersed and do not form preferential interactions. (B) Fraction of helix-helix contacts ($f_{\text{helix-helix}}$) for each residue of WT, G335A, G338A, A326P, and M337P in blue, orange, green, red and purple, respectively. (C) Correlation between the average residue helicity and fraction of involvement in helix-helix contacts for WT, G335A, G338A, A326P, and M337P in blue, orange, green, red and purple, respectively. Dashed lines are the linear fit with $R^2$ shown in the key.

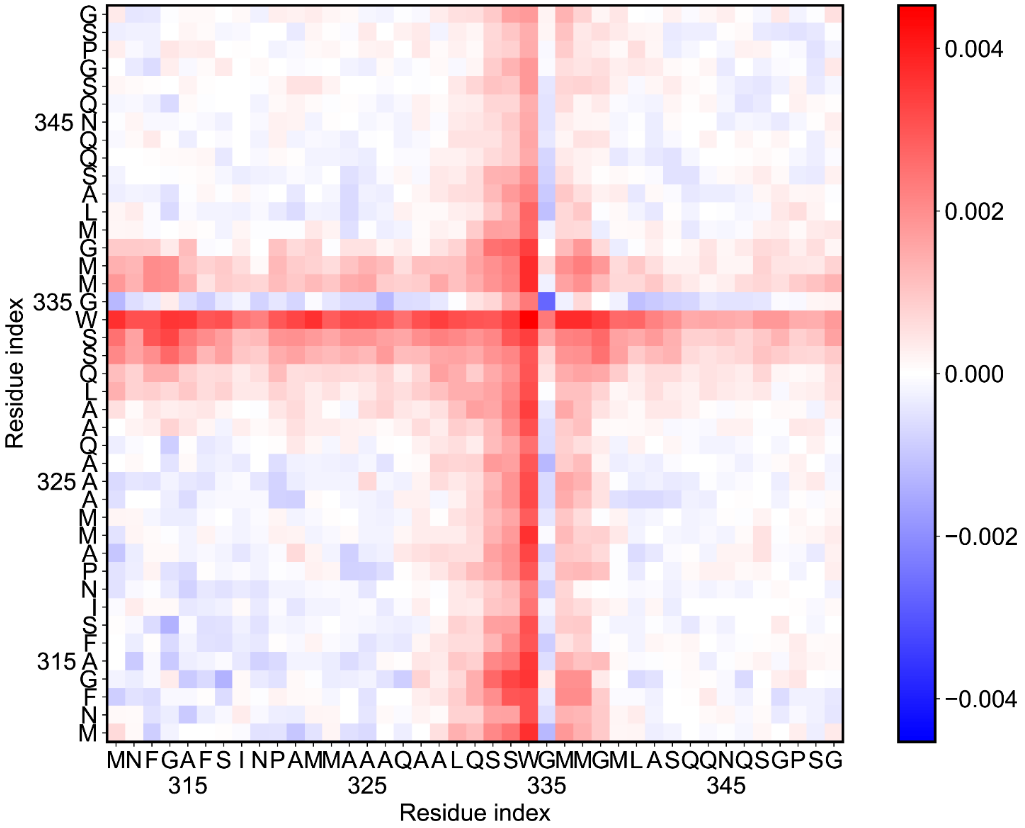


**Figure S11.** Probability differences of intermolecular backbone-backbone contacts between WT and $WT^{+\Delta hel}$ TDP-43 CR condensates The difference was calculated through subtracting the contact probabilities in the condensed phase of WT by those of $WT^{+\Delta hel}$. Positive values (red) represent higher contact probabilities in the WT case, while negative values (blue) represent lower contact probabilities compared to the $WT^{+\Delta hel}$ case.

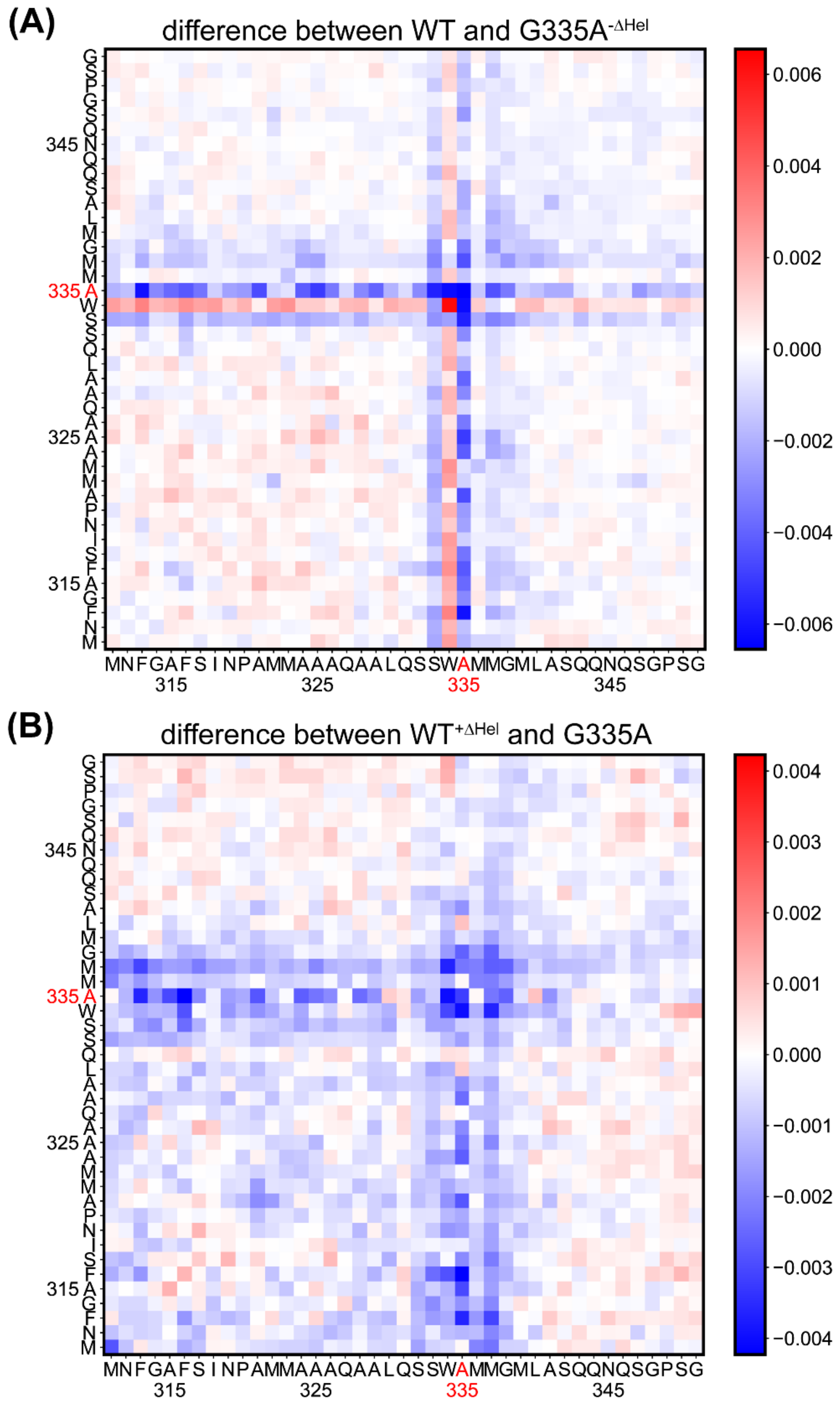


**Figure S12.** The effect of Ala on intermolecular residue-residue contact probabilities. (A-B) Probability differences of inter-molecular residue-residue contacts between WT and G335A$^{-\Delta hel}$ and between WT$^{+\Delta hel}$ and G335A. The difference was calculated through subtracting the contact probabilities in the condensed phase of former variant by those of the latter one. Positive values (red) represent higher contact probability in the former variant, while negative values (blue) show lower contacts compared to the latter one.

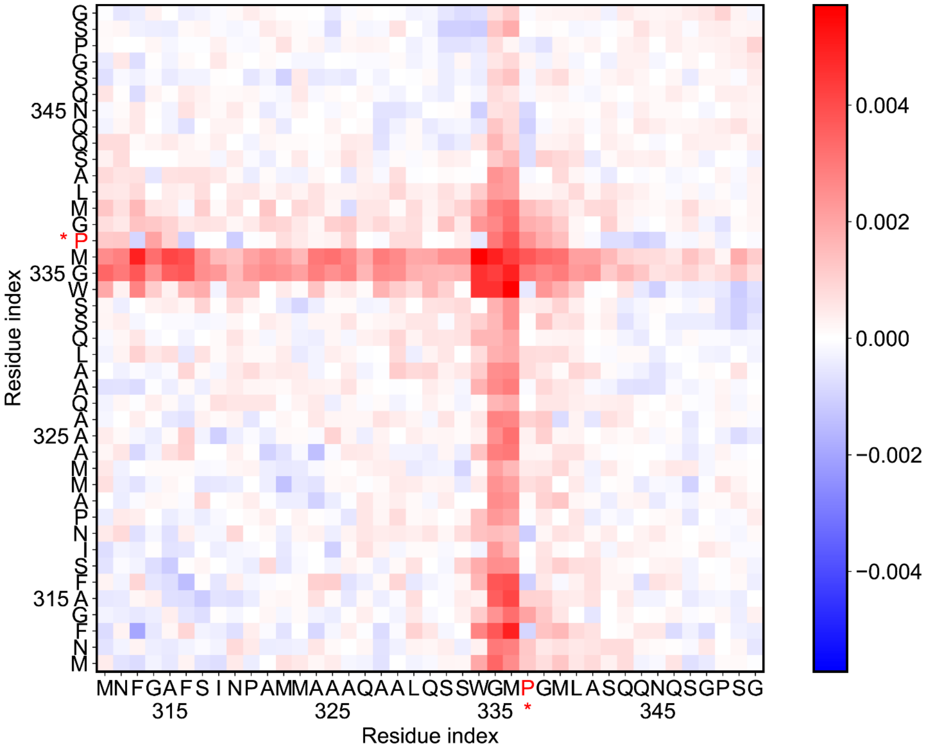


**Figure S13.** Probability differences of inter-molecular residue-residue contacts between WT and M337P. The difference was calculated through subtracting the contact probabilities in the condensed phase of WT by that of M337P. Positive values (red) represent higher contact probability in the WT case, while negative values (blue) show lower contact probabilities compared to the M337P case.

(13) Schuster, B. S.; Dignon, G. L.; Tang, W. S.; Kelley, F. M.; Ranganath, A. K.; Jahnke, C. N.; Simpkins, A. G.; Regy, R. M.; Hammer, D. A.; Good, M. C.; et al. Identifying sequence

perturbations to an intrinsically disordered protein that determine its phase-separation behavior. *P Natl Acad Sci USA* **2020**, *117* (21), 11421-11431. DOI: 10.1073/pnas.2000223117.
(14) Bremer, A.; Farag, M.; Borcherds, W. M.; Peran, I.; Martin, E. W.; Pappu, R. V.; Mittag, T. Deciphering how naturally occurring sequence features impact the phase behaviours of disordered prion-like domains. *Nat Chem* **2022**, *14* (2), 196-+. DOI: 10.1038/s41557-021-00840-w.
(15) Conicella, A. E.; Dignon, G. L.; Zerze, G. H.; Schmidt, H. B.; D'Ordine, A. M.; Kim, Y. C.; Rohatgi, R.; Ayala, Y. M.; Mittal, J.; Fawzi, N. L. TDP-43 alpha-helical structure tunes liquid-liquid phase separation and function. *P Natl Acad Sci USA* **2020**, *117* (11), 5883-5894. DOI: 10.1073/pnas.1912055117.
(16) Brady, J. P.; Farber, P. J.; Sekhar, A.; Lin, Y. H.; Huang, R.; Bah, A.; Nott, T. J.; Chan, H. S.; Baldwin, A. J.; Forman-Kay, J. D.; et al. Structural and hydrodynamic properties of an intrinsically disordered region of a germ cell-specific protein on phase separation. *P Natl Acad Sci USA* **2017**, *114* (39), E8194-E8203. DOI: 10.1073/pnas.1706197114.
(17) Conicella, A. E.; Dignon, G. L.; Zerze, G. H.; Schmidt, H. B.; D’Ordine, A. M.; Kim, Y. C.; Rohatgi, R.; Ayala, Y. M.; Mittal, J.; Fawzi, N. L. TDP-43 α-helical structure tunes liquid–liquid phase separation and function. *Proceedings of the National Academy of Sciences* **2020**, *117* (11), 5883-5894. DOI: doi:10.1073/pnas.1912055117.
